# Systematic mining and functional analysis of factors regulating wheat spike development for breeding selection

**DOI:** 10.1101/2022.11.11.516122

**Authors:** Xuelei Lin, Yongxin Xu, Dongzhi Wang, Yiman Yang, Xiaoyu Zhang, Xiaomin Bie, Zhongxu Chen, Yiliang Ding, Long Mao, Xueyong Zhang, Fei Lu, Xiansheng Zhang, Cristobal Uauy, Xiangdong Fu, Jun Xiao

**Author notes:** These authors contributed equally to this article (Xuelei Lin, Yongxin Xu, Dongzhi Wang). Correspondence: Jun Xiao.

## Abstract

The spike architecture of wheat plays a crucial role in determining grain number, making it a key trait to optimize in wheat breeding programs. In this study, through a multi-omic approach, we analyzed the transcriptome and epigenome profiles of the shoot apex at eight developmental stages, revealing coordinated changes in chromatin accessibility and H3K27me3 abundance during the flowering transition. We constructed a core transcriptional regulatory network (TRN) that drives wheat spike formation, and experimentally validated a multi-layer regulatory module involving TaSPL15, TaAGLG1, and TaFUL2. By integrating the TRN with genome-wide association analysis (GWAS), we identified 227 transcription factors (TFs), including 42 with known functions and 185 with unknown functions. Further investigation of 61 novel TFs using multiple homozygous mutant lines uncovered 36 TFs with altered spike architecture or flowering time, such as TaMYC2-A1, TaMYB30-A1, and TaWRKY37-A1. Of particular interest, TaMYB30-A1, downstream and repressed by WFZP, was found to regulate fertile spikelet number. Notably, during the domestication and breeding process in China, the excellent haplotype of *TaMYB30-A1* containing a C allele at the WFZP binding site was enriched, leading to improved agronomic traits. Our study presents novel and high-confidence regulators and offers an effective strategy for understanding the genetic basis of wheat spike development, with practical impact for wheat breeding applications.

## Introduction

Bread wheat (*Triticum aestivum* L.) is a globally important crop, contributing to >20% of human caloric and protein consumption (Giraldo et al., 2019; Xiao et al., 2022). Grain yield is determined by the architecture of the wheat inflorescence which exhibits a distinct compound spike morphology with multiple sessile spikelets (Gauley and Boden, 2019; Koppolu et al., 2022). The development of the wheat spike involves several transitions, starting from the shoot apical meristem (SAM) producing leaf, node, and internode primordia, transitioning to the inflorescence meristem (IM) which elongates and forms a double-ridged structure. The upper ridge gives rise to the spikelet meristem (SM), while the lower ridge undergoes arrested leaf primordia development. Successively, the SM gives rise to the glume and lemma primordia, followed by the appearance of the floret primordium. In the apex, the IM transitions to a terminal spikelet that determines the maximum spikelet number of the spike. This stage is accompanied by stamen primordia emergence and floral organ differentiation. Pistils and awns subsequently elongate, resulting in indeterminate spikelets that when mature will typically containing 4-6 fertile florets each (Waddington et al., 1983; Vahamidis et al., 2014). It is the spikelet arrangement (spikelet number per spike, SNS) together with the spikelet fertility which determines the grain number per spike (GNPS) and ultimately grain yield in wheat (Boden et al., 2015; Sakuma and Schnurbusch, 2020; Luo et al., 2023).

Recent studies have demonstrated that modifying spike architecture can substantially enhance grain yield in wheat, underscoring the importance of this trait (Wang et al., 2022; Zhang et al., 2022). Significant advancements have been made in unraveling the molecular regulation of cereal inflorescence architecture, particularly in wheat (Gao et al., 2019; Gauley and Boden, 2019; Koppolu and Schnurbusch, 2019; Yuan et al., 2020; Koppolu et al., 2022; Luo et al., 2023). *PHOTOPERIOD-1* (*PPD1*), *FLOWERING LOCUS T1* (*FT1*), and *FT2* coordinately control the transition and duration of spike development (Boden et al., 2015; Finnegan et al., 2018; Shaw et al., 2019). MADS-box transcription factors (TFs) VERNALIZATION1 (VRN1) and its paralogs FRUITFULL2 (FUL2) and FUL3 are necessary for SM identity (Li et al., 2019a; Li et al., 2021). Wheat TEOSINTE BRANCHED1 (TaTB1) interacts with FT1 to control the production of paired spikelets (Dixon et al., 2018). *WHEAT ORTHOLOG OF APO1* (*WAPO1*) affects spikelet number, while *Q* (*APETALA 2-like gene 5*, *AP2L5*) reduce it when inactive by regulating the formation of terminal spikelet (Kuzay et al., 2019; Debernardi et al., 2020). *WFZP*, a homolog of rice *FRIZZY PANICLE* (*FZP*), influences spikelet number and identity (Li et al., 2021; Du et al., 2021). Additionally, *SQUAMOSA promoter-binding protein-like 13-2B* (*TaSPL13-2B*), *TaSPL14*, and *AGAMOUS-LIKE6* (*TaAGL6*) are involved in determining spikelet number (Li et al., 2020; Cao et al., 2021; Kong et al., 2022). However, functional genomics research in wheat is still limited due to challenges in gene identification methods.

Advancements in sequencing technologies have greatly contributed to the field of population genetics (Sehgal and Dreisigacker, 2022). Large-scale genome-wide association studies (GWAS) utilizing single nucleotide polymorphism (SNP) have been instrumental in identifying genetic loci associated with spike developmental traits in wheat, including spike length (SL), SNS, GNPS, grain setting, and spike compactness (Hao et al., 2020; Pang et al., 2020; Li et al., 2022). However, GWAS alone provides limited insights into causal variants and genes due to linkage disequilibrium. To address this, gene co-expression networks have been utilized, leveraging stage-specific transcriptome data and population-wide RNA-seq to identify key regulators (Wang et al., 2017; Li et al., 2018; VanGessel et al., 2022). Moreover, investigations into chromatin features have revealed associations with tissue-specific or stress-induced gene expression and subgenome-biased expression patterns in wheat (Li et al., 2019b; Concia et al., 2020; Yuan et al., 2022; Zhao et al., 2023). Integration of gene expression, chromatin features, and transcription factor binding sites has enabled the prediction of gene regulatory networks, as demonstrated in recent studies (Chen et al., 2023; Tang et al., 2023). By combining meta agronomic traits GWAS analysis with gene regulatory networks, the identification of candidate genes associated with relevant regions can be enhanced (Chen et al., 2023). However, the lack of chromatin landscape data from tissues during wheat spike development has hindered the exploration of spike-specific gene regulatory networks, limiting the discovery of novel regulators for spike architecture.

In this study, we aimed to enhance our understanding of the interplay between chromatin landscape and transcriptome dynamics during wheat spike formation. To achieve this, we conducted a comprehensive analysis of the transcriptome and epigenome profiles of the developing spike of the elite wheat cultivar Kenong 9204 (KN9204) across different developmental stages. By integrating co-expression and regulatory networks specific to spike development with GWAS, we identified key factors influencing wheat spike architecture. Furthermore, we validated multiple novel genes and conducted in-depth analyses to unravel the potential functions of these regulators and elucidate the allelic selection processes that occur during the breeding process.

## Results

### A transcriptome and chromatin landscape atlas for wheat spike development

To understand the transcriptional regulation for wheat spike development, we sampled eight distinct developmental stages of winter wheat cultivar KN9204, including SAM, transition apex (W1.5), early double ridge stage (W2), double ridge stage (W2.5), glume primordium stage (W3), lemma primordium stage (W3.25), floret primordium stage (W3.5), and late terminal spikelet stage (W4) (**Figure 1A**, developmental stages as per Waddington et al., 1983). RNA-seq and the Assay for Transposase Accessible Chromatin with high-throughput sequencing (ATAC-seq) were used to profile global gene expression and chromatin accessibility, respectively (**Figure 1A and Supplemental Figure 1A**) (Buenrostro et al., 2015; Kaya-Okur et al., 2019; Zhao et al., 2023).

**Figure 1.**
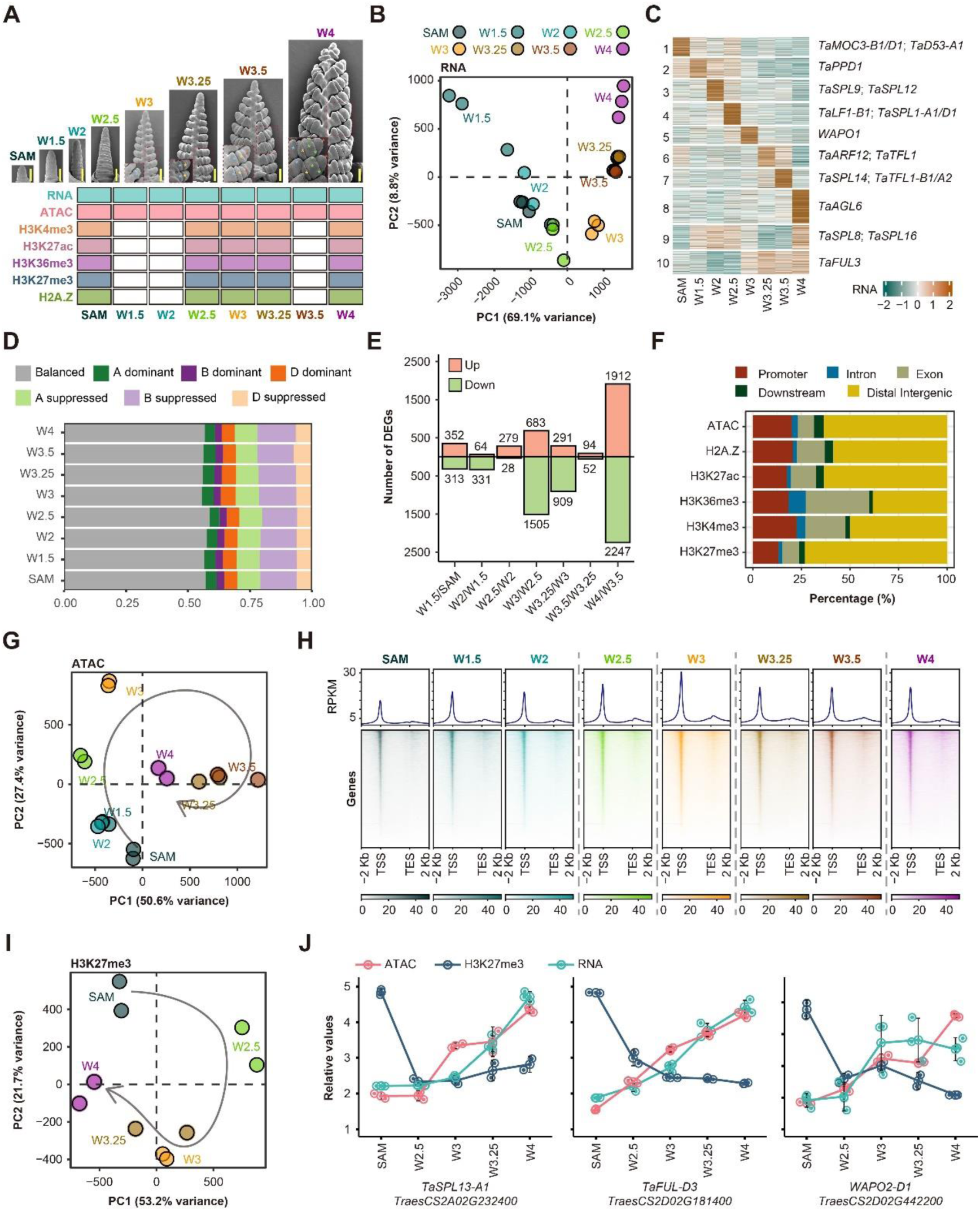
Transcriptome and chromatin landscapes during wheat spike development. **A,** Scanning electronic micrographs (SEM) of young spikes in eight developmental stages and experimental design. Scale bars = 200 μm. **B,** Principal component analysis (PCA) of transcriptome showing distinct developmental stages. Each sample is represented by a dot, with spike development stages in different colors. Three biological replicates were sequenced for each stage. **C,** Heatmap of expressed genes sorted by k-means clustering across the samples collected at different developmental stages. Representative genes of each cluster are listed. **D,** Homoeolog expression bias of differentially expressed triads across spike developmental stages. **E,** Number of DEGs between adjacent developmental stages. DEGs were defined by DESeq2 with the threshold of |log2 (Fold Change)| ≥ 1 and FDR ≤ 0.05. **F,** Peak distribution of ATAC-seq and various histone modifications relative to genes. **G,** PCA of ATAC-seq samples during spike development. Each dot represents one sample and two biological replicates were sequenced for each stage. Color represents different stages and the curved arrow shows the developmental trajectory. **H,** Chromatin accessibility dynamics in proximal regions (promoter and genic regions) during wheat spike development. RPKM was used for data normalization. **I,** PCA of H3K27me3 samples during spike development. Each dot represents one sample. Two biological replicates were sequenced for each stage. **J,** Gene expression, chromatin accessibility and H3K27me3 changes of representative genes during spike development. The Y-axis indicates relative values of Z-scaled gene expression, chromatin accessibility and H3K27me3 levels.

A total of 58,875 high-confidence genes were identified, expressing in at least one stage (TPM > 0.5) (**Supplemental Table 1**). The stages were divided into two groups: the vegetative to flowering transition group (SAM, W1.5, W2, W2.5) and the reproductive growth phase group (W3, W3.25, W3.5, W4) (**Figure 1B and Supplemental Figure 1B**). We found stage-specific expressed genes through clustering (**Figure 1C**), including known factors involved in flowering time regulation and inflorescence development. *PPD1*, with *Ppd-A1b/Ppd-B1a/Ppd-D1a* alleles in KN9204 (Wu et al., 2021), is responsible for flowering time and expressed before the transition to flowering. (Beales et al., 2007; Boden et al., 2015; Perez-Gianmarco et al., 2020), is expressed before the transition to flowering. *WAPO1*, involved in maintaining SM activity (Kuzay et al., 2022), shows high expression at W3 stage. *TaTFL1*, correlated with SNS (Wang et al., 2017; Sun et al., 2023), is highly expressed at W3.25 and W3.5 stages, while *TaAGL6*, a floral organ regulator (Kong et al., 2022), is highly expressed at W4 stage (**Figure 1C**). We further analyzed the expression pattern of homoeolog triads given the hexaploid nature of bread wheat (Ramírez-González et al., 2018). The majority of triads (68.2% to 69.5% across stages) were categorized as balanced, with single-homoeolog suppressed triads being more common than single-homoeolog dominance triads (**Supplemental Figures 1C and 1D**). B-homoeolog dominance was significantly more frequent than A- or D-homoeolog dominance (**Supplemental Figure 1D**). A similar pattern was observed in DEGs triads, but with fewer balanced triads (55.6%-58.7%) and a higher proportion of dominant (12.0%-13.6%) and suppressed (29.3%-31.2%) triads (**Figure 1D**). Morphological transition points, such as W2.5 to W3, W3 to W3.25, and W3.5 to W4, showed more DEGs between adjacent stages, coinciding with glume primordium, lemma primordium, and stamen primordium initiation, respectively (**Figure 1E and Supplemental Table 2**).

Similar to previous studies (Zhao et al., 2023), we found that accessible chromatin in wheat is mainly located in the distal intergenic regions and promoters (**Figure 1F**). Chromatin accessibility around the transcriptional start site (TSS) positively correlates with gene expression (**Supplemental Figure 1E**). During wheat spike development, there is a roughly continuous trajectory of chromatin accessibility changes, with stages grouped into five sub-clusters (**Figure 1G and Supplemental Figure 1F**). Chromatin accessibility increases from vegetative to flowering transition and inflorescence initiation, then declines (**Figure 1H and Supplemental Figure 1G**). The fraction of triads with non-balanced chromatin accessibility increased first and then decreased, with a higher proportion of non-balanced chromatin accessibility during the flowering transition (**Supplemental Figure 1H**).

For histone modification profiling, we examined SAM, W2.5, W3, W3.25, and W4 stages based on the chromatin accessibility sub-clusters via Cleavage Under Targets and Tagmentation (CUT&Tag) (**Figure 1A**). H3K27ac, H3K4me3, and H3K36me3 around TSS positively correlate with gene expression, while H3K27me3 is enriched in the gene body of no/low-expressed genes (**Figure 1F and Supplemental Figure 1E**). H3K27me3 and H3K36me3 are mutually exclusive, while H3K4me3 and H3K27ac show a positive correlation (**Supplemental Figure 1I)**. Various histone modifications exhibit stage-specific transitions during developmental stages (**Figure 1I and Supplemental Figures 1J**). H3K27me3 has a continuous trajectory and plays a significant role in regulating gene expression (**Figure 1J**). Dominant expressed homoeologs have higher chromatin accessibility, active histone marks, and lower repressive histone marks, while suppressed homoeologs show the opposite pattern (**Supplemental Figure 2A**). B-subgenome histone marks are generally lower than those of A- and D-subgenomes (**Supplemental Figure 2B**). Overall, we have generated transcriptomic and epigenomic profiles of wheat spike development stages, shedding light on chromatin regulation during this process.

### Permissive chromatin status facilitates vegetative-to-reproductive transition

During the transition from the vegetative to reproductive stage, there was a global increase in chromatin accessibility (**Figure 1H**). We specifically examined differentially accessible promoter regions (DAPRs) at various stages, including SAM, W1.5, W2, W2.5, and W3. A total of 49,153 DAPRs were identified and grouped into six clusters (**Figure 2A and Supplemental Table 3**). Most of the DAPRs showed increased accessibility at either the W2.5 or W3 stage, including clusters 2, 3, 4, and 6 (**Figure 2A and Supplemental Table 4**). Interestingly, genes in these clusters exhibited a strong positive correlation between chromatin accessibility and transcriptional changes (*R* > 0.5) (**Figure 2B**). We observed that the chromatin accessibility shifted from balanced to suppressed at W2-W2.5 and W2.5-W3 stages for some triads, and then transitioned back to balanced at W3-W3.25 (**Supplemental Figures 3A and 3B**). Specifically, genes with increased chromatin accessibility at the W2.5 or W3 stage significantly overlapped with genes showing elevated expression levels compared to SAM (1,920 genes in gene set I; **Figure 2C, Supplemental Figure 3C and Supplemental Table 5**). This synchronous increase in chromatin accessibility and gene expression highlights their close association (**Figure 2D**). The degree of increased chromatin accessibility is highly correlated (*R* = 0.85) with the fold change in elevated expression levels (**Figure 2E**). Gene Ontology (GO) analysis revealed enrichment in hormone biosynthesis and signaling, inflorescence meristem identity, and asymmetric cell division in gene set I (**Figure 2F**).

**Figure 2.**
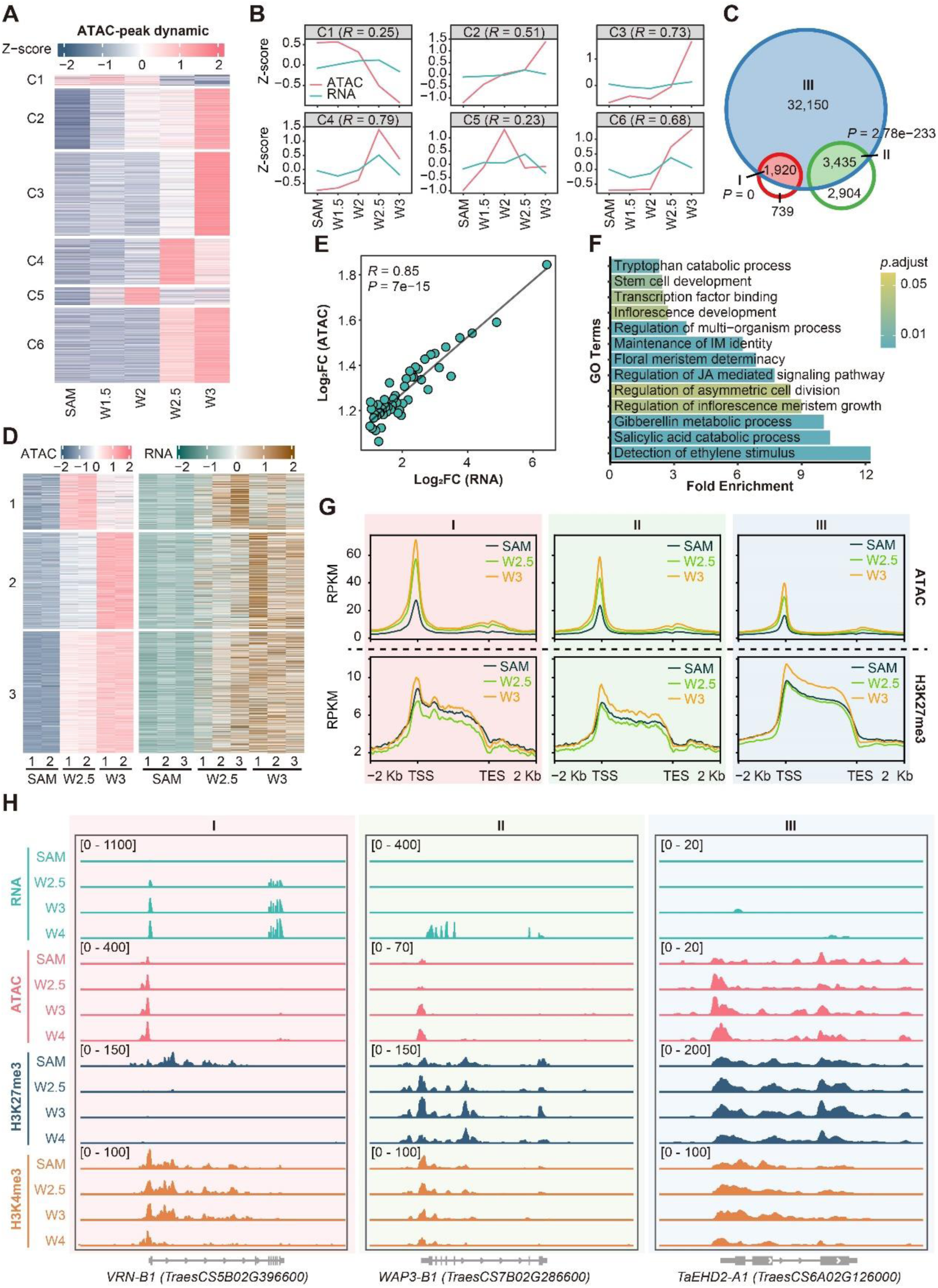
Chromatin accessibility and H3K27me3 dynamics facilitate vegetative-to-reproductive transition. **A,** K-means clustering of differentially accessible promoter regions (DAPR) across the samples collected from SAM to W3 stage. **B,** The chromatin accessibility and gene expression tendency for each cluster in (**A**). *R* value stands for the Pearson correlation coefficients. **C,** Overlapping of gene groups. Blue circle, genes with gained-chromatin accessibility of cluster 2, 3, 4, 6 in (**A**); Red circle, genes elevated expression at W2.5 and W3 stage versus SAM stage; Green circle, genes up-regulated at W3.25, W3.5, W4 stage versus former stages. Gene set I: overlap between red and blue circle genes; gene set II: overlap between red and green circle; gene set III: remaining genes from blue circle. Hypergeometric test was used to calculate *P* value for the enrichment of gene set I and II. **D,** Synchronous pattern of gene set I between the gain of chromatin accessibility and elevation of gene expression from SAM to W2.5 and W3 stage. Biological replicates are shown separately. **E,** Pearson correlation of gene expression elevation (X-axis) and chromatin accessibility gain (Y-axis) from SAM to W2.5 or W3 stage based on 1,920 genes in gene set I in (**C**). Genes were ranked by fold change of elevated expression and separated into 50 bins. *P* values are determined by the two-sided Pearson correlation coefficient analysis **F,** GO enrichment analysis of 1,920 genes in gene set I. **G,** Chromatin accessibility (top) and H3K27me3 (bottom) levels of genes in gene set I, II, III at SAM, W2.5 and W3 stages. **H,** Genomic tracks showing expression, chromatin accessibility and histone modifications change at representative genes of *VRN-B1* (gene set I, left), *WAP3-B1* (gene set II, middle) and *TaEHD2-A1* (gene set III, right).

**Figure 3.**
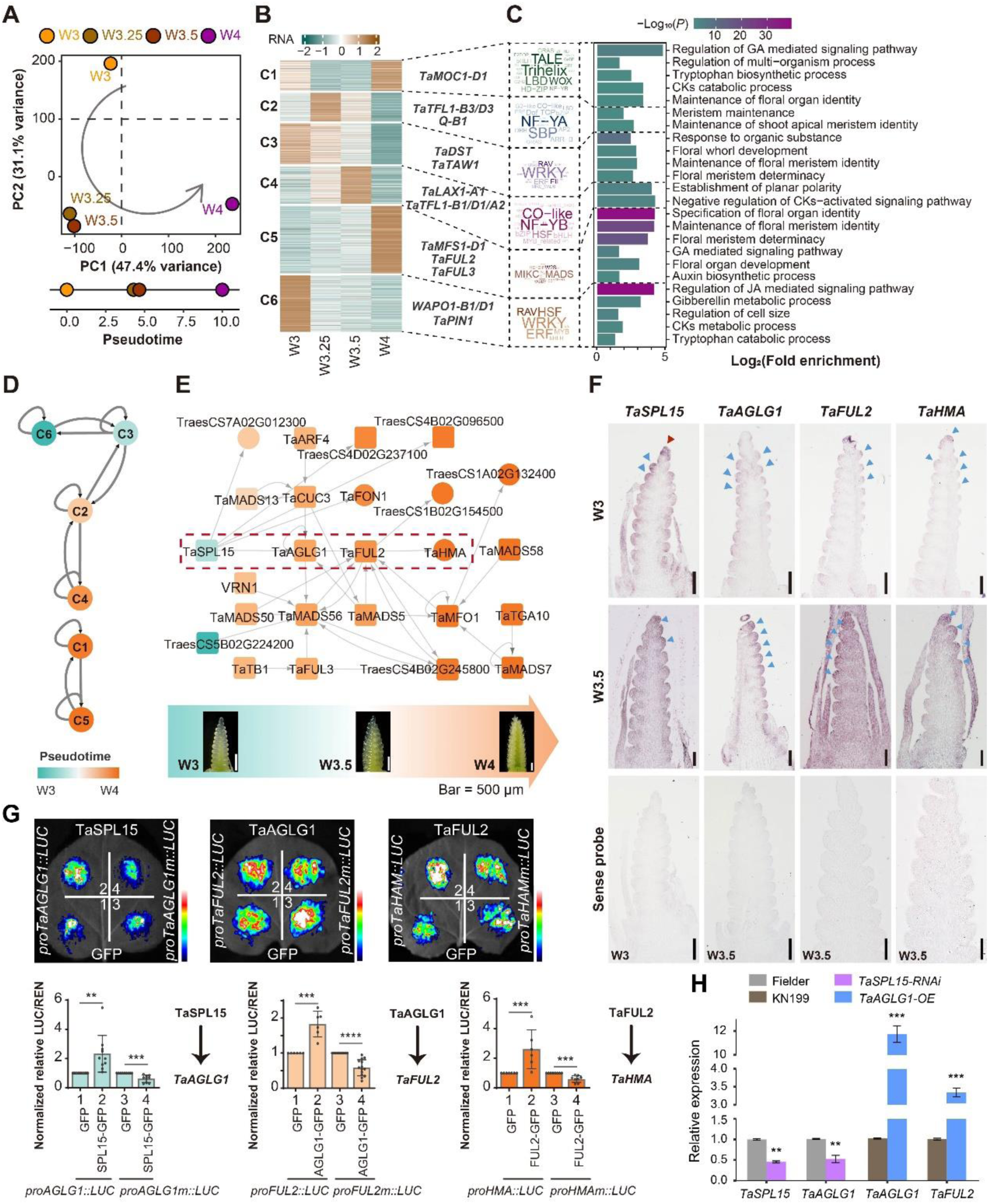
Transcriptional regulatory network (TRN) governing spike architecture formation. **A,** PCA plots of gene expression from W3 to W4 stages. The developmental time units (DTU) values calculated based on the scaled straight distance between two adjacent stages are indicated in the lower panel. **B,** K-means clustering of DEGs from W3 to W4 stages and representative genes in each cluster. **C,** TF family enrichment and GO enrichment analysis within each cluster in (**B**). **D,** Hierarchical transcriptional regulations of sequentially expressed gene clusters from (**B**). **E,** TRN for represented key TFs participated in spike development, from W3 to W4 stages. Genes were organized by the expression timing from left to right as indicated by different color gradients. TFs and non-TF coding genes are represented by rectangles or circles, respectively. The dashed red rectangle frame indicates a four-layer regulation module. **F,** Spatiotemporal expression patterns of *TaSPL15*, *TaAGLG1*, *TaFUL2* and *TaHMA* at different spike developmental stages, as indicated by *in situ* hybridization. Sense probes of each gene were detected in each stage as negative controls. The red and blue triangles indicate gene expression position (see text). Scale bars = 100 μm. **G,** Luciferase reporter assay validation of transcriptional regulation among representative TF-target pairs as indicated in the red rectangle frame of (**E**). Mutations of the TF binding sites were introduced in the promoter region of target genes separately. Student’s *t* test was used for the statistical significance. **, *P* ≤ 0.01; ***, *P* ≤ 0.001. **H,** The expression level of *TaSPL15* and *TaAGLG1* in Fielder and *TaSPL15*-*RNAi* transgenic plants, and *TaFUL2* and *TaAGLG1* in KN199 and *TaAGLG1-OE* transgenic plants, estimated by qRT-PCR using young spike at W3.5 stage. Expression level of genes in Fielder and KN199 is set as 1.0 separately, and the relative expression of each gene in transgenic plants is shown as average ±SD of three replicates. Student’s *t* test was used for the statistical significance. **, *P* ≤ 0.01; ***, *P* ≤ 0.001.

Furthermore, open chromatin can establish a "primed" state for future gene activation (Bonifer and Cockerill, 2017; He and Li, 2018). We identified 3,435 genes with gained chromatin accessibility at the W2.5 or W3 stage but elevated mRNA levels at stages after W3 (gene set II; **Figure 2C, Supplemental Figure 3C and Supplemental Table 5**). These genes are primarily involved in hormone metabolism, floral organ identity, and meristem development (**Supplemental Figure 3D**). In contrast, most of genes (85.7%) gained chromatin accessibility but without consequent changes in mRNA levels at the sampled spike developmental stages (gene set III; **Figure 2C and Supplemental Table 5**). These genes are associated with polarity specification, cell size regulation, and the regulation of translation or protein transport (**Supplemental Figure 3E**). The different expression patterns of gene sets gained chromatin accessibility suggesting that open chromatin alone is not sufficient to activate gene expression.

In addition, the histone modification H3K27me3 showed a close association with the transcriptional patterns of gene sets I, II, and III (**Figure 2G**). Conversely, other histone modifications such as H3K27ac, H3K4me3, and H2A.Z did not exhibit significant correlations (**Supplemental Figure 3F**). Gene set I displayed higher chromatin accessibility at the SAM stage, with a notable increase at W2.5 and W3 stages, accompanied by a decrease in H3K27me3 levels at W2.5 (**Figure 2G**). The example of the flowering promoting factor VRN1 (*VRN-B1* locus for representation) demonstrated increased chromatin accessibility in the proximal promoter and decreased H3K27me3 in both the promoter and first intron regions (**Figure 2H**), which are crucial for regulating *VRN1* transcription (Yan et al., 2004; Fu et al., 2005). In gene set II, although chromatin accessibility was elevated at W3 and W3.25 stages, the relative openness remained low, and there was no significant change in the H3K27me3 level (**Figure 2H**). This combined effect suggests a "primed" transcriptional status, where gene expression increases later with the reduction of H3K27me3, as observed in the wheat *APETALA3-B1* (*WAP3-B1*) gene (Hama et al., 2004) (**Figures 2G and 2H**). Gene set III exhibited the lowest chromatin accessibility and the highest coverage of H3K27me3, which restricts gene activation, as observed in the *EH-domain-containing protein 2* (*TaEHD2-A1*) gene (**Figures 2G and 2H**).

### Transcriptional regulatory network for spike architecture formation

After flowering transition (up to W2.5), inflorescences undergo several developmental stage transitions that eventually lead to the terminal spikelet stage (W4) which defines the final SNS, contributing to grain yield (**Figure 1A and Figure 3A**). By calculating the pseudo time based on PCA distance (**Figure 3A**), we obtained a developmental trajectory from W3 to W4, with W3.25 and W3.5 stages being closely related. A total of 8,200 DEGs were identified and categorized into six clusters (**Figure 3B and Supplemental Table 6**). Genes related to floral meristem determinacy and hormone metabolism were highly expressed at W3 stage, while hormone signaling and polarity establishment genes were active at W3.25 and W3.5 stages. At the W4 stage, genes associated with floral organ identity and polarity specification were elevated (**Figures 3B and 3C**). Among the stage-specific DEGs, we focused on enriched TFs. ERF and WRKY TFs were enriched and highly expressed at the W3 stage within clusters 3 and 6. NF-Y and SBP TFs were abundantly expressed at the W3.25 and W3.5 stages within clusters 2 and 4, respectively. MADS-box TFs stood out at the W4 stage within cluster 5 (**Figures 3B and 3C**).

Accessible chromatin regions serve as binding sites for TFs. Based on the recognition of TF-motifs and the co-expression patterns of TFs and their downstream targets, we constructed a TRN by integrating the ATAC-seq and RNA-seq time course data (**Supplemental Figure 4A**). A total of 36,908 pairs of 5,106 TF-target gene interactions were identified, with 4,916 pairs involving regulation among TFs (**Supplemental Figure 4B**). To facilitate visualization and exploration of the TRN, we constructed a website accessible to the public (http://39.98.48.156:8800/#/, see methods for details). The TRN reveals a clear sequential regulatory relation among gene clusters expressed from the W3 to W3.5 stages, namely C6-C3-C2-C4 (**Figure 3D**). Additionally, the W4 stage-specific expressed genes in clusters C1 and C5 are separate and regulated independently (**Figure 3D**), indicating distinct regulatory networks for spikelet and floret development. We predicted high number of targets for members of the ERF, B3, TCP, DOF, and MIKC-MADS TF families, supporting their core regulatory roles (**Supplemental Figure 4C**). Notably, we identified functionally characterized factors involved in wheat spike development, such as TaTB1 (Dixon et al., 2018), VRN1, and TaFUL2, TaFUL3 (Li et al., 2019a; Li et al., 2021), in the network (**Figure 3E**).

To evaluate the predictive power of the TRN, we extracted a module that contains factors that have been individually studied but without known regulatory relationships. This module includes TaSPL15 (Pei et al., 2023), AGAMOUS-LIKE GENE1/PANICLE PHYTOMER2 (TaAGLG1/PAP2) (Yan et al., 2003, Wang et al., 2017), and TaFUL2 (Li et al., 2019a; Li et al., 2021) (**Figure 3E**). We performed *in situ* hybridization to characterize the spatial-temporal expression profile of these factors (**Figure 3F and Supplemental Figure 4H**). *TaSPL15* was strongly expressed in the IM (red triangle shown at W3 stage) and spikelet meristem (blue triangles shown at W3 and W3.5 stage). *TaAGLG1* and *TaFUL2* were highly expressed in the spikelet meristem (blue triangles), whereas *heavy metal ATPase* (*TaHMA*), a potential downstream target of TaFUL2, was also expressed in the spikelet meristem (blue triangles) (**Figure 3F and Supplemental Figure 4H**). The temporal expression pattern and the presence of specific motifs in the open chromatin regions suggest a positive regulatory hierarchy involving the TaSPL15-TaAGLG1-TaFUL2-TaHMA module (**Supplemental Figures 4D and 4E**). Indeed, we observed a positive transcriptional regulatory circuit among them using a luciferase (LUC) reporter assay in tobacco leaves. This regulation is dependent on the specific recognition of TFs and cis motifs (**Figure 3G and Supplemental Figure 3F**). Consistent with the *in vitro* LUC reporter assay (**Figure 3G**), *TaAGLG1* was significantly down-regulated in *TaSPL15-RNAi* wheat (**Figure 3H and Supplemental Figure 3G)**, and *TaFUL2* was up-regulated in *TaAGLG1-OE* plants (**Figure 3H**). Thus, the constructed TRN effectively predicted regulatory relationships between TFs during spike development.

**Figure 4.**
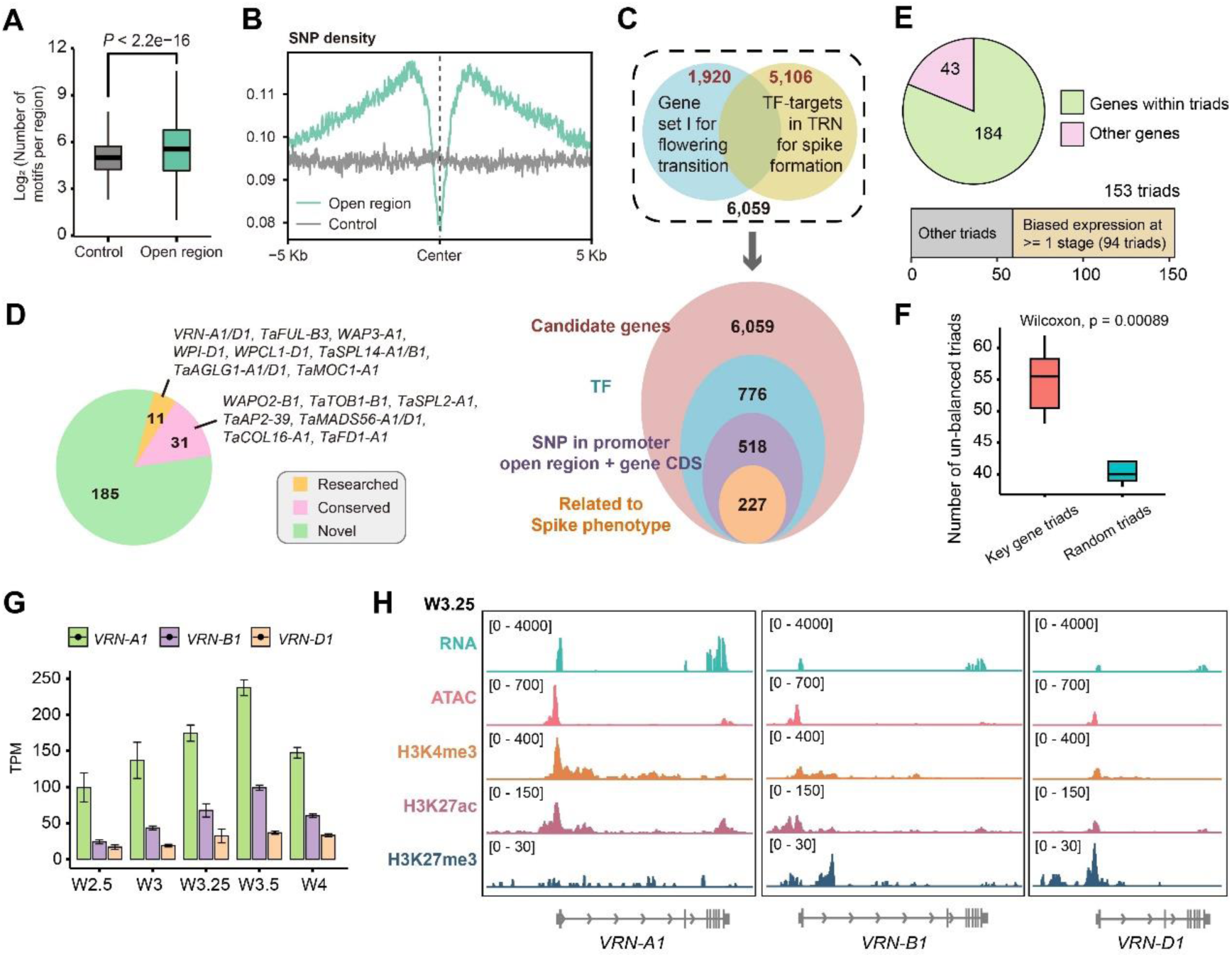
Screening for key wheat spike developmental regulators by integration of multi dimensional data. **A.** Enrichment of TF binding motifs in open chromatin regions. Green and gray refer to open chromatin regions and control regions, respectively. Intergenic closed chromatin regions with the same number and size as the open chromatin region were randomly selected as control. The bars within box plots represent 25^th^ percentiles, medians, and 75^th^ percentiles and Wilcoxon rank sum test was used to determine significant differences. **B.** SNP density in 5 Kb regions flanking open chromatin regions (green) and control regions (gray). Open chromatin regions show low sequence diversity in contrast to the control. **C.** Schematic of the strategy used for identification of wheat spike architecture TFs. **D.** Proportion of candidate genes for spike development in each category as indicated. Representative genes are listed. **E.** The summary of gene expression of sub-genomes homeologs for the 227 selected TFs. The pie chart displays the number of TFs with or without triads. For the TFs with triads, the proportion of triads with balanced (gray square) or unbalanced (yellow square) expressed patterns are shown in the bottom panel. Triads with unbalanced expression at any stages were defined as unbalanced triads. **F.** The proportion of unbalanced expressed triads for the 153 key TF triads (red) in (**E**) and randomly selected triads (green) across eight stages. The bars within box plots represent 25^th^ percentiles, medians, and 75^th^ percentiles. Wilcoxon rank sum test was used to determine significant differences. **G.** Expression level of *VRN1* homoeologs across five development stages. **H.** Genomic tracks showing expression, chromatin accessibility and histone modifications of *VRN1* homoeologs at W3.25 stage.

### Integration with population genetics to screen regulators shaping wheat spike

Meta GWAS studies have successfully identified genetic loci associated with spike developmental traits in wheat (Hao et al., 2020; Pang et al., 2020; Abou-Elwafa and Shehzad, 2021). To further investigate regulators that govern spike architecture, we combined publicly available GWAS data (**Supplemental Table 7**) with key regulators identified for floral transition (**Figure 2**) and factors within the transcriptional regulatory network (TRN) specifically generated for spike formation (**Figure 3**). In addition to gene coding sequence variations, we recognized the importance of open chromatin regions in establishing transcriptional regulatory circuits (Klemm et al., 2019). Notably, open chromatin regions were found to have significantly more TF recognition motifs (**Figure 4A**). Moreover, TF binding footprints within these open chromatin regions exhibited lower DNA variation frequency in 172 common wheat accessions (Zhou et al., 2020) compared to flanking regions and random genomic backgrounds, indicating their functional significance (**Figure 4B**).

Starting with 1,920 genes from gene set I for flowering transition (**Figure 2C**) and 5,106 TFs-targets within the TRN (**Supplemental Figure 3B**), we focused on TF coding genes and identified 776 candidate TFs. Among them, more than 65% (518 TFs) showed DNA variations in the promoter open chromatin region or coding region with non-synonymous mutations (Zhou et al., 2020; Hao et al., 2020). Eventually, 227 TFs were found to harbor DNA variations significantly associated with spike related traits within 2 Mb regions centered around GWAS signals (**Figure 4C and Supplemental Table 8**). Notably, eleven factors among these TFs have already been functionally studied in wheat (**Figure 4D and Supplemental Table 9**), including *VRN-A1/D1* (Li et al., 2019a), *TaFUL-B3* (Li et al., 2019a), *WAP3-A1* (Hama et al., 2004). Additionally, 31 factors are considered ’conserved’ orthologues, studied for inflorescence development in other species, such as *WAPO2-B1* (*ABERRANT PANICLE ORGANIZATION 2*, *APO2*, Ikeda-Kawakatsu et al., 2012), *TaTOB1-B1* (*TONGARI-BOUSHI1*, *TOB1*, Tanaka et al., 2012) and *TaSPL2-A1* (*SPL2*, Wang et al., 2016) (**Figure 4D and Supplemental Table 9**). Intriguingly, the majority of the identified candidate TFs (n = 185) have unknown functions in wheat and other crops and were thus considered ’novel’ (**Figure 4D**). These TFs are enriched for ERF, MYB, WRKY, bHLH, and B3 TF families (**Supplemental Table 9**). Approximately one-fifth of the 227 TFs are not founds as triads of homoeologous genes, and 61.4% (94/153) of those present as triads showed unbalanced expression in one or more stages (**Figure 4E**), a significantly higher proportion than control triads (**Figure 4F**). For example, *VRN-A1* exhibited higher expression than *VRN-B1* and *VRN-D1* (**Figure 4G**). Notably, *VRN-D1* was suppressed with repressive chromatin status, showing low chromatin accessibility and less active histone modifications (H3K4me3, H3K27ac) but heavy repressive histone modification H3K27me3 (**Figure 4H**).

In summary, the integration of GWAS and TRN analyses identified hundreds of TFs potentially involved in spike development, with many being novel and showing unbalanced expression patterns. This suggests that manipulating a single homoeologue may lead to phenotypic changes in wheat spike architecture.

### Identification of novel regulators for spike development

To evaluate the effectiveness of our strategy in identifying novel regulators for wheat spike development, we conducted a comprehensive investigation of spike developmental defects using indexed chemical-induced (TILLING) mutant lines in both hexaploid wheat Cadenza (Krasileva et al., 2017) and KN9204 (Wang et al., 2023). Among the 185 novel TFs, a total of 122 TFs were found to have at least one KN9204 mutant line containing a loss-of-function mutation (**Figure 5A and Supplemental Table 9**). To ensure robust results, we further narrowed down the candidates by looking for TFs with loss-of-function mutations in multiple mutant lines, with at least one homozygous line, resulting in the identification of 61 TFs that met these criteria. Out of them, 36 TFs (∼ 59%) exhibited notable spike developmental defects that could be categorized into three main types: differences in flowering time (Type-I, n = 11); degeneration of spikelet or floret (Type-II, n = 19); alterations in spikelet number or spikelet density (SD) (Type-III, n = 17) (**Figures 5B and 5C and Supplemental Table 9**).

**Figure 5.**
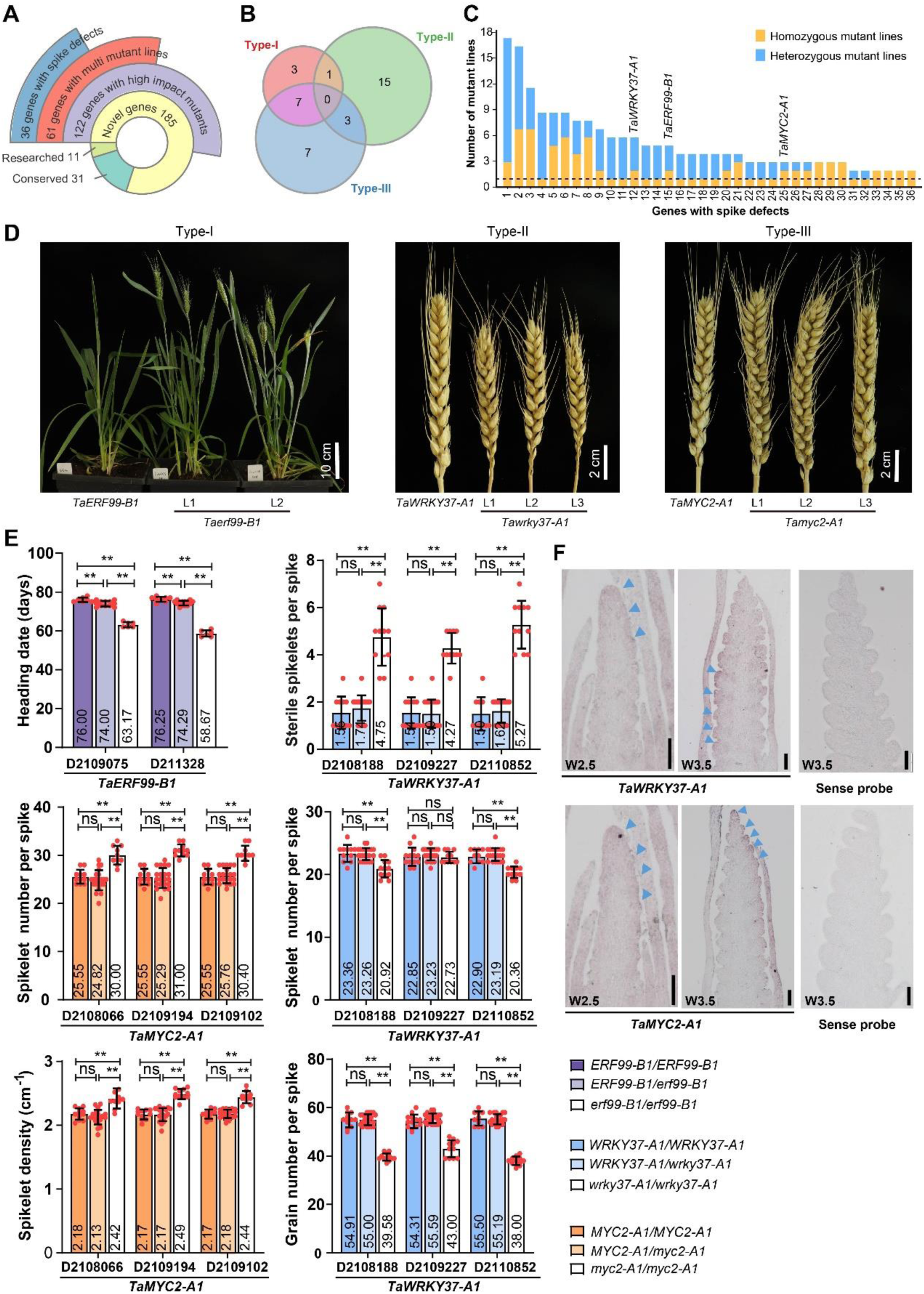
Novel transcription factors identified by TRN regulate wheat spike development. **A,** Summary of novel TFs with different categories of KN9204 TILLING mutant lines. **B,** Venn-diagram of different types of spike-related phenotype present in KN9204 TILLING mutant lines containing a homozygous mutation at novel TFs coding genes. Type-I, flowering time difference; Type-II, degeneration of spikelet or floret; Type-III, SD or SNS difference. **C,** Overview of 36 novel TFs with spike-related phenotypes based on the number of homozygous (orange) and heterozygous (blue) mutant lines. Horizontal dashed line indicates TFs with multiple mutant lines. The three novel TFs used for phenotypic analyses are labeled. **D,** Representative KN9204 TILLING mutant lines of type-I/II/III spike developmental defects as compared to control KN9204. L1, L2 and L3 represent different homozygous mutant lines. **E,** Quantification of different types of spike developmental defects in KN9204 TILLING mutant lines. Heading date of *Taerf99-B1*; SSPS, SL, SNS and GNPS of *Tawrky37-A1*; SNS and SD of *Tamyc2-A1*. Phenotypes of the BC_2_F_2_ homozygous mutant (white), heterozygous (light color), and wild-type (dark color) sibling plants. The error bars denote ±SD, and each plant is represented as a dot. Student’s *t*-test was used for the statistical significance. *, *P* ≤ 0.05; **, *P* ≤ 0.01; ns, no significant difference. **F,** Spatiotemporal expression patterns of *TaWRKY37-A1* and *TaMYC2-A1* at W2.5 and W3.5 stages, as indicated by *in situ* hybridization. The blue triangles indicate gene expression position (see text). Sense probe of each gene was detected as negative control. Scale bars = 100 μm.

To firmly establish the causal relationship between the identified TFs and the observed spike developmental defects, we conducted backcrossing experiments for 30 mutant lines, covering a subset of ten genes. Among them, 28 out of 30 backcrossing populations show expected co-segregation of the spike development defects (**Supplemental Table 10**). We successfully identified representative isogenic sibling lines in the backcross population for TFs exhibiting type-I, type-II, and type-III phenotypes (**Figure 5D**). Mutant lines of TraesCS1B02G282300 (*TaERF99-B1*, type-I) displayed a two-week early flowering phenotype compared to that of the wild-type sibling lines. Mutant lines of TraesCS2A02G443800 (*TaWRKY37-A1*, type-II) had an increase in sterile spikelets per spike (SSPS) at the basal part of the inflorescence, leading to a decrease in SNS and GNPS compared to their wildtype counterparts. Furthermore, the mutant lines of TraesCS1A02G193200 (*TaMYC2-A1*, type-III) showed increased SD, SNS and GNPS compared to their wild type control plants (**Figures 5D and 5E and Supplemental Figure 5A**). It is noteworthy that similar phenotypes were also observed in independent Cadenza knock-out mutant lines for *TaWRKY37-A1* and *TaMYC2-A1* (**Supplemental Figures 5B and 5C**). Interestingly, *TaERF99* and *TaWRKY37* exhibit B-homoeolog suppression and D-homoeolog dominance, respectively (**Supplemental Figure 5D**), supporting the results that single homoeolog mutants lead to phenotypic consequences. We further examined the spatiotemporal expression pattern of those genes by *in situ* hybridization (**Figure 5F**). Consistent with their morphologic defects, *TaWRKY37-A1* and *TaMYC2-A1* are both expressed in the SM initiation region (blue triangles at W2.5 stage). However, their expression diverges at W3.5 stage; while *TaWRKY37-A1* is highly expressed in the SM at the base of spike (consistent with rudimentary basal spikelets), *TaMYC2-A1* is highly expressed in the spikelet meristem at the upper region of the spike (**Figure 5F**). These results exemplify the validity of the approach and successfully demonstrate the significant roles played by numerous novel TFs identified through our integration strategy in regulating spike development.

### Regulation of wheat spike architecture by novel factor TaMYB30-A1

To further explore the application of integrated TRN and GWAS in dissecting the molecular function of individual genes in wheat spike development, we took a novel regulator identified in the TRN (TraesCS6A02G224200; *TaMYB30-A1*) for in-depth examination (**Supplemental Figure 4B**). *TaMYB30-A1* was located within a genetic region that showed significant association with SSPS based on GWAS analysis (**Supplemental Figure 6A and Supplemental Table 11**). The peak SNP (chr6A:421426337; G/A) showed significant differences in SSPS and explained 4.8% of SSPS variation in the population (**Supplemental Figure 6A and Supplemental Table 11**). *TaMYB30-A1* lacks the B genome homoeolog, whereas the D homoeolog exhibits low expression levels during spike development, potentially attributed to relatively low chromatin accessibility and high H3K27me3 levels (**Supplemental Figures 6B and 6C**). *TaMYB30-A1* shows distinct expression patterns during spike development based on *in situ* hybridization. It is first detected at the SM initiation region at W2.5, and then it exhibits high expression in the spikelet and floral meristem during the W3.5, W4, and W4.25 stages (**Figure 6A**, blue triangles). During floret organ development from W4 to W4.5 (with the carpel primordium present), *TaMYB30-A1* is detected in the stamen (green stars), pistil primordium (red triangle), and carpel primordium (yellow star) (**Figure 6A**). These expression patterns are consistent with its putative role in regulating spikelet number and floret development.

**Figure 6.**
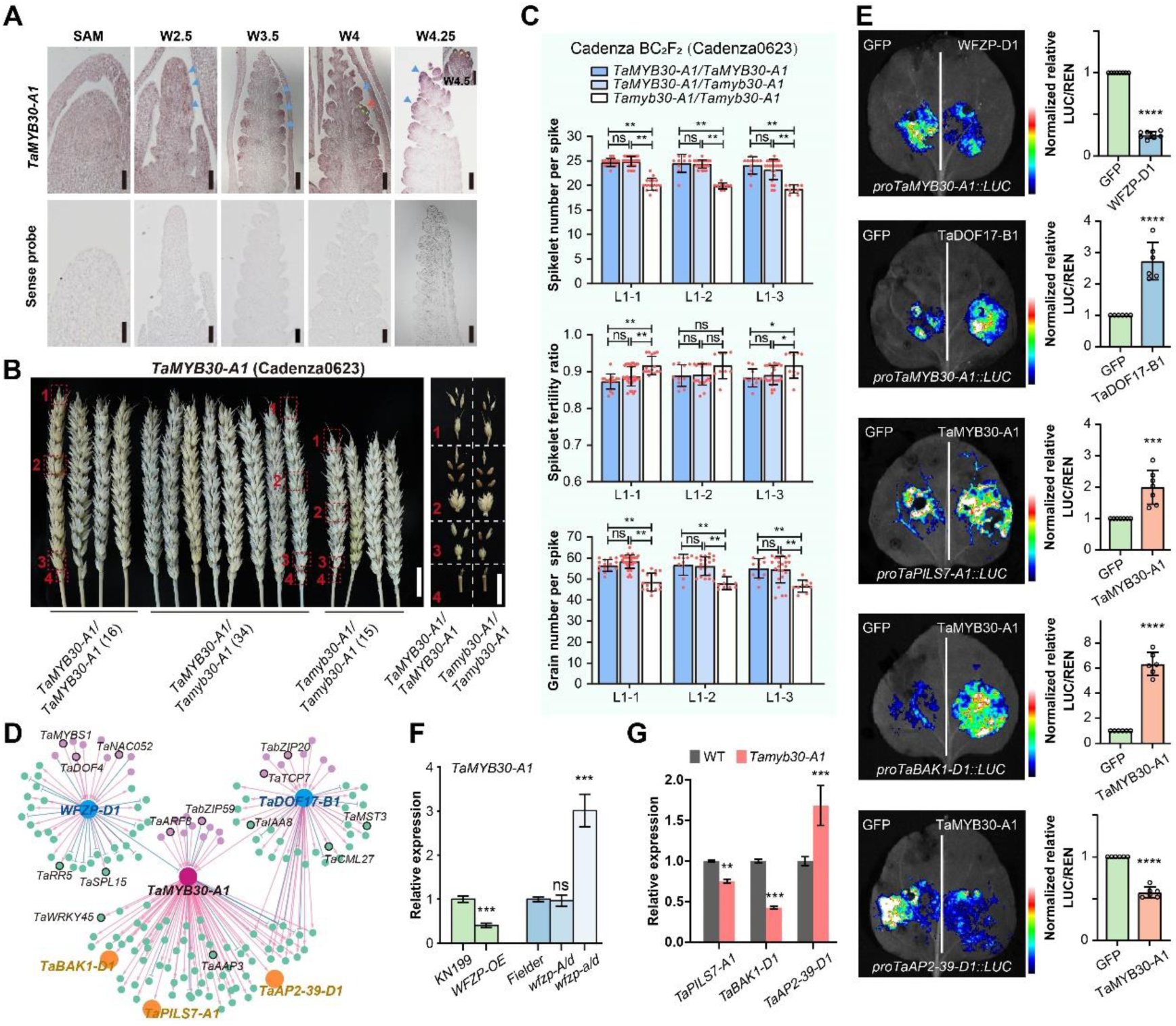
*TaMYB30-A1* regulates spike architecture. **A,** Spatiotemporal expression pattern of *TaMYB30-A1* is indicated by *in situ* hybridization. The blue/red triangles and green/yellow stars indicate gene expression position (see text). Sense probe was detected as negative control. Scale bars = 100 μm. **B,** The spike phenotype of *Tamyb30-A1* mutant plants (Cadenza0623). The *Tamyb30- A1* mutants were backcrossed with wild-type Cadenza, and the BC_2_F_2_ were used to perform the co-segregation test. Spikelets corresponding to the numbers and positions (red dotted box) were shown on the right for fertility. Scale bars = 2 cm. **C,** Statistical comparison of SNS, spikelet fertility ratio and GNPS between Cadenza and *Tamyb30-A1* mutant lines. L1-1, L1-2 and L1-3 represent independent sibling plants from the same mutant line Cadenza0623. The error bars denote ± SD, SNS and spikelet fertility ratio of each plant are presented as dots. Student’s *t*-test was used for the statistical significance. *, *P* ≤ 0.05; **, *P* ≤ 0.01; ns, no significant difference. **D,** The TRN of *TaMYB30-A1*. The upstream and downstream genes verified using luciferase reporter assays were highlighted in blue and orange, respectively. **E,** Luciferase reporter assays of TaMYB30-A1 regulatory network. Student’s *t*-test was used for the statistical significance. ***, *P* ≤ 0.001; ****, *P* ≤ 0.0001. **F,** The expression level of *TaMYB30-A1* at W3.5 stage in KN199, *WFZP*-OE transgenic plant and Fielder, *wfzp-A/d*, *wfzp-a/d* CRISPR lines by RT-qPCR. The error bars denote ±SD. Student’s *t*-test was used for the statistical significance. ***, *P* ≤ 0.001; ns, no significant difference. **G**, Expression level of *TaMYB30-A1* target genes in *Tamyb30-A1* mutant at W3.5 stage. **, *P* ≤ 0.01; ***, *P* ≤ 0.001.

We acquired loss-of-function mutant lines of *TaMYB30-A1* from both KN9204 and Cadenza mutant populations, which carry premature termination codons at positions 108 and 89, respectively (**Figure 6B and Supplemental Figures 6D and 6E**). To reduce mutation load, we performed backcrossing with wild-type plants and evaluated the genotype-phenotype co-segregation (**Figure 6B and Supplemental Figures 6D-6F**). The homozygous *Tamyb30-A1* mutant plants displayed significant reductions in SL and SNS, but an increase in spikelet fertility ratio, compared to segregated heterozygous and wild-type plants in the BC_2_F_2_ population (**Figures 6B and 6C and Supplemental Figure 7A**). Notably, the total GNPS was significantly lower in *Tamyb30-A1* mutants compared to controls, mainly due to a significant reduction in SNS (∼10%; *P* < 0.01) (**Figure 6C**). Similar trends were observed in the co-segregation test of the KN9204 BC_1_F_2_ population for SL, SNS, and spikelet fertility ratio, although the reduction in GNPS was not as prominent (**Supplemental Figures 6D and 7B**).

**Figure 7.**
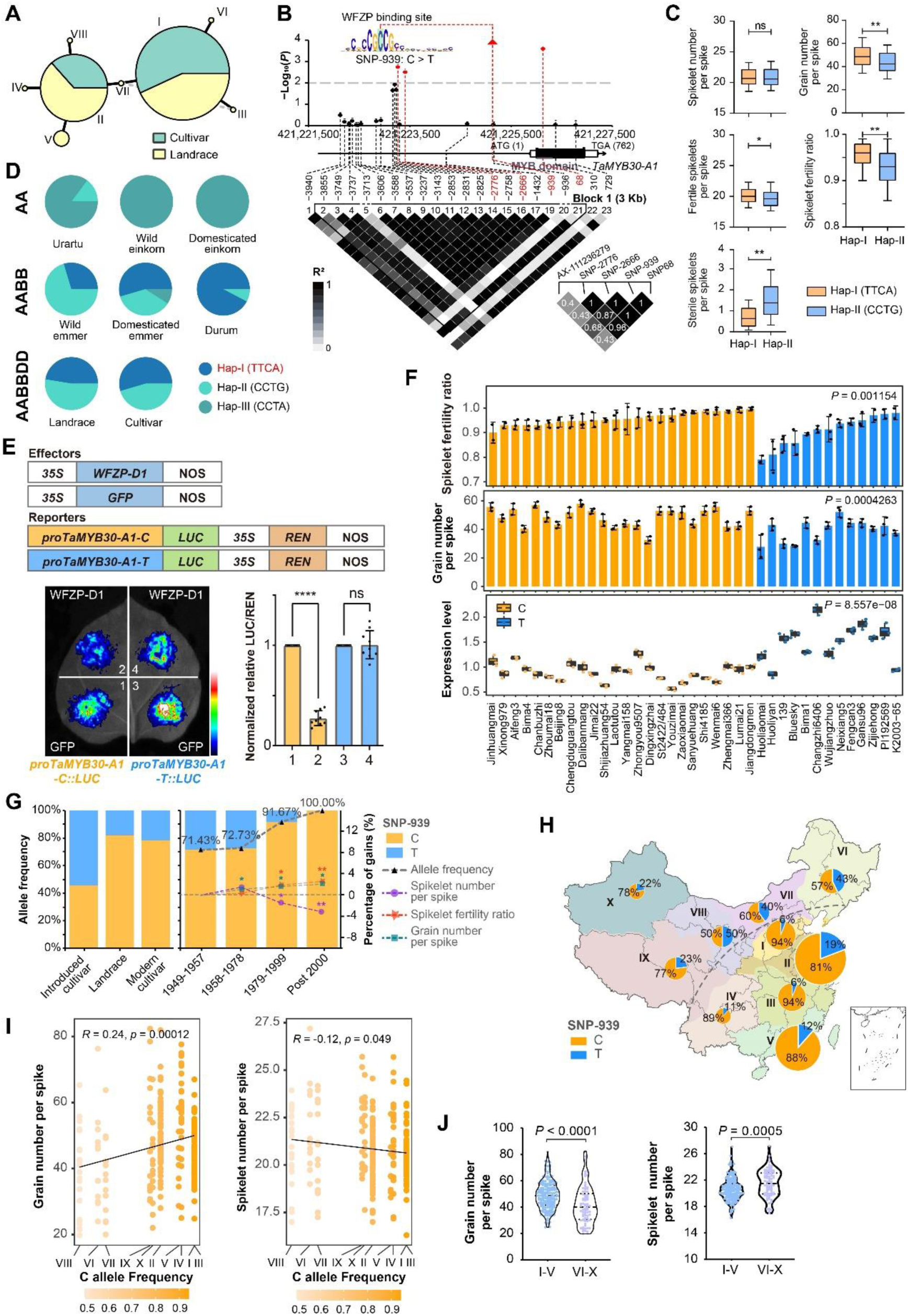
*TaMYB30-A1* haplotype underwent selection during domestication and breeding in China. **A,** Haplotype network of *TaMYB30-A1* representing two major haplotypes. Circle size is proportional to the number of samples within a given haplotype, and dashes between haplotypes represent unobserved, inferred haplotypes. Lines between haplotypes represent mutational steps between alleles. Green, cultivars; Yellow, landrace. **B**, Association analysis between genetic variations in *TaMYB30-A1* and SSPS. The schematic diagram of the ∼6 Kb genomic region of *TaMYB30-A1* is shown, followed by a LD plot with white to black representing *r^2^* ranging from 0 to 1. Dots connected with the red dashed lines indicate significant variants (-log_10_(*P*) =2), with the most significantly associated SNP (SNP -939) marked by a red triangle. The linkage disequilibrium of these four SNPs with GWAS peak signal AX-111236279 were displayed in the lower right of the figure. SNP -939 was located in the core WFZP DNA-binding motif. **C**, Comparison of SNS, spikelet fertility ratio and GNPS between haplotypes defined by four significantly associated SNPs. Wilcoxon rank-sum test was used to determine the statistical significance; *, *P* < 0.05; **, *P* < 0.01. **D**, Frequency of *TaMYB30-A1* haplotypes in different species. Hap-I (TTCA) was considered as excellent haplotype and marked in red. **E**, Luciferase reporter assays of *TaMYB30-A1* promoter activity with the WFZP binding site C/T mutation. Schematic diagram showing the vectors used. Student’s *t*-test was used for the statistical significance; ****, *P* < 0.0001; ns, no significant difference. **F,** The spikelet fertility ratio, GNPS and *TaMYB30-A1* expression level of 37 cultivars with different *TaMYB30-A1* haplotypes. The expression level of *TaMYB30-A1* was determined by RT-qPCR using young spike at W3.5 stage, with three biological replicates for each cultivar. Cultivars with C-allele and T-type were marked in yellow and blue, respectively. The spikelet fertility ratio and GNPS are shown with data from each environment as a dot. Significant difference between the two haplotypes was determined by Student’s *t*-test. **G,** The percentage of accessions with reference and alternative allele of SNP -939 in different categories (left) and breeding periods (right). Lines indicate the allele frequency of reference C-allele (black, triangle), SNS (purple, dot), spikelet fertility ratio (red, star), and GNPS (blue, square), respectively. Wilcoxon rank-sum test was used to determine the statistical significance with accessions 1949-1957. Only those that reached the significance threshold are labeled (with the color corresponding to the lines). *, *P* < 0.05; **, *P* < 0.01. **H,** The percentage of accessions with reference and alternative allele of SNP -939 in different ecological zones of China. The size of pie charts in the geographical map shows the number of cultivars, with percentages of the two SNP haplotypes in different colors (C-allele, yellow; T-allele, blue). A gray dotted line was used to distinguish the zones I-V (with frequency of C-type more than 80%) and VI-X (with frequency of C-allele less than 80%). **I,** The distribution of GNPS and SNS in ten ecological zones and their correlation with the frequency of C allele at SNP -939 of *TaMYB30-A1*. From left to right, the C allele frequency in ecological zones increases. The GNPS is positively while SNS is negatively correlative with the C allele frequency with the Pearson correlation analysis. **J,** Zones I-V have lower SNS but higher GNPS than zones VI-X. Wilcoxon rank-sum test was used to determine the statistical significance between two groups.

We identified a transcription regulatory module containing *TaMYB30-A1* in the TRN (**Figure 6D**). Among the potential upstream regulators, *WFZP-D1* and TraesCS2B02G592700 (*TaDOF17-B1*) were found to either repress or activate *TaMYB30-A1*, respectively, in reporter assays conducted in tobacco leaves (**Figure 6E and Supplemental Figure 7C**). These results are consistent with the temporal expression patterns of *WFZP-D1*, *TaDOF17-B1*, and *TaMYB30-A1* during spike development. *TaDOF17-B1* is specifically up-regulated at W2.5 coinciding with the increase in *TaMYB30-A1* expression. Analogously, *WFZP-D1* expression increased at W3.25 stage coinciding with *TaMYB30-A1* reduction (**Supplemental Figure 7D**). The repression of *TaMYB30-A1* by WFZP was further validated through the analysis of transgenic lines. Consistent with our model, over-expression of *WFZP* and CRISPR-induced *wfzp-a/d* mutations (Li et al., 2021; Du et al., 2021), led to significant down-and up-regulation of *TaMYB30-A1*, respectively (**Figure 6F**). Among the numerous potential downstream targets of TaMYB30-A1 (**Figure 6D**), several genes, including *TraesCS2D02G425700* (*APETALA-2-Like Transcription Factor 39*, *TaAP2-39-D1*), *TraesCS7D02G416900* (*Brassinosteroid insensitive 1-associated kinase 1*, *TaBAK1- D1*), and *TraesCS5A02G354300* (*PIN-LIKES 7*, *TaPILS7-A1*) were selected since their orthologs are known to regulate inflorescence development in rice (Yaish et al., 2010; Khew et al., 2015; Yi et al., 2021). TaMYB30-A1 activated *TaBAK1-D1* and *TaPILS7- A1*, and repressed *TaAP2-39-D1* in reporter assays conducted in tobacco leaves (**Figure 6E**). Again, these results are consistent with their temporal expression pattern during spike development (**Supplemental Figure 7D**). In addition, the expression of *TaBAK1- D1* and *TaPILS7-A1* was up-regulated, while *TaAP2-39-D1* was down-regulated, in the *Tamyb30-A1* homozygous mutant lines of the Cadenza BC_2_F_2_ population (**Figure 6G**), providing further evidence for TaMYB30-A1 regulation of these targets.

### Selection of *TaMYB30-A1* excellent haplotype during domestication and breeding

Next, we explored the effect of different *TaMYB30-A1* haplotypes on spike architecture. We identified eight haplotypes in the common wheat population (Guo et al. 2020; Hao et al.2020; Zhou et al., 2020), with Hap-I being most common in cultivars whereas Hap-II was more prevalent in landraces (**Figure 7A**). Among the DNA variations across the *TaMYB30-A1* locus (**Supplemental Figure 8A**), four SNPs in high linkage disequilibrium with GWAS peak signal AX-111236279 were significantly associated with SSPS, and accessions with the Chinese Spring allele (Hap-I, TTCA) showed superior agronomic characteristics, with lower SSPS while higher FSPS, spikelet fertility ratio and GNPS compared to Hap-II CCTG, thus Hap-I was defined as the excellent haplotype (**Figures 7B and 7C and Supplemental Figure 8B**). We further analyzed the origin of Hap-I in the *Triticum* population and found that this haplotype emerged in tetraploid wheat and underwent human selection during the domestication of both tetraploid and hexaploid wheat (**Figure 7D and Supplemental Figure 8C**).

The most significant DNA variation of Hap-I/II associated with SSPS was SNP -939 (C/T), located in the *TaMYB30-A1* promoter region and within the WFZP core binding motif (**Figures 7B and Supplemental Figure 8B**). We hypothesized that this SNP could influence WFZP-mediated repression of *TaMYB30-A1* and hence affect its expression. Indeed, in a reporter assay in tobacco leaves, WFZP repressed *TaMYB30- A1*, and this repression was abolished with the C-to-T mutation equivalent to SNP -939 (**Figure 7E**). Among the 37 wheat accessions from the GWAS panel (24 C : 13 T), those with the C allele showed a higher ratio of fertile spikelets (more GNPS) and a significantly lower level of *TaMYB30-A1* expression compared to T allele (**Figure 7F and Supplemental Figure 8D**). This data support the model in which the T variation at position -939 increases *TaMYB30-A1* expression via a reduction in WFZP repression and is consequently associating with reduced spikelet fertility but without significant reduction on spikelet number.

In the Chinese wheat mini-core collection, the percentage of accessions carrying the C-allele was higher in landraces (82.1%) and modern cultivars (78.2%) compared to introduced cultivars (45.5%) (**Figure 7G, left**). The frequency of the C-allele steadily increased with cultivar year of release during wheat breeding in China. Along with this increase in frequency, the spikelet fertility ratio and GNPS steadily increased, while SNS showed a decrease (**Figure 7G, right**). Interestingly, the alleles showed distinct distribution characteristics in the major agro-ecological zones of China (**Figure 7H**). The frequency of the C-allele showed a positive correlation with GNPS, while displaying a negative correlation with SNS across diverse agro-ecological zones (**Figure 7I**). Regions I-V, characterized by a relatively higher C-allele frequency, exhibited higher GNPS and spikelet fertility ratio, but lower average SNS or shorter SL. Conversely, regions VI-X, with lower C-allele frequency, showed the opposite trend (Figure 7J and Supplemental Figure 8E).

In conclusion, the excellent Hap-I (TTCA) of *TaMYB30-A1* likely first appeared in tetraploid wheat, undergoing strong human selection during domestication in both tetraploid and hexaploid wheat. The SNP -939 influence the WFZP-mediated repression of *TaMYB30-A1*, and has been strongly selected during the breeding process in China over the past 40 years. It is tempting to speculate that this could be due to the need to balance spikelet number and fertility to adapt to different agro-ecological zones (**Supplemental Figure 8F**).

## Discussion

Spikelet and floret development play a crucial role in shaping inflorescence architecture and ultimately determining grain productivity in wheat. In this study, we conducted a comprehensive profiling of the epigenomic landscape over time, including various histone modifications, chromatin accessibility, and transcriptomes of developing wheat spikes, spanning from vegetative development (SAM) to terminal spikelet stage where spike architecture is determined. This enabled us to analyze the transcriptional regulation governing the critical processes of flowering transition, spikelet formation, and floret development, all of which significantly impact wheat inflorescence architecture (Sakuma and Schnurbusch, 2020; VanGessel et al., 2022; Luo et al., 2023).

### Chromatin control of vegetative-to-reproductive transition in wheat

The shoot apical meristem is responsible for generating primordia that initiate different tissues, such as leaf primordia during the vegetative stage and spikelet primordia as the plant transitions to flowering. These processes are tightly regulated by various transcription factors such as TaTB1, VRN1, TaFT1, and TaPPD1, responding to internal and external signals (Koppolu and Schnurbusch, 2019; Luo et al., 2023). Gene expression and cell identity are closely intertwined with chromatin status (Perino and Veenstra, 2016). During the transition from SAM to W2.5/W3 stage (vegetative-to-reproductive growth), chromatin accessibility increases. However, not all open chromatin leads to gene expression changes, especially when the genic region is covered by H3K27me3. Yet, the chromatin accessibility gains are synchronized with the activation of genes during floral induction, including those involved in inflorescence meristem identity and hormone biosynthesis and signaling. Moreover, open chromatin can create a primed state for genes to activate at a later developmental stage when H3K27me3 is removed. Notably, flowering time gene *VRN-B1* (Shimada et al., 2009) and *TaEHD2-A1* (Matsubara et al., 2008), as well as genes associated with spikelet meristem formation like *WAP3-B1* (Hama et al., 2004), are influenced by chromatin status. Thus, a permissive chromatin environment is crucial for activating key regulators in initiating reproductive development in wheat.

### Systematic and efficient identification of novel regulators for spike development

Cell identity transition, driven by hormone signaling and transcriptional regulatory networks, plays a crucial role in the initiation, distribution, and termination of spikelets (Youssef and Hansson, 2019; VanGessel et al., 2022; Qi et al., 2019). Understanding these regulatory networks can offer potential candidates for shaping inflorescence structure. Genetic variations in key regulatory factors or changes in the regulatory circuit can influence inflorescence architecture outcomes. Leveraging our time-series profiling of transcriptome and chromatin accessibility data, along with TF-motif binding information from model plants (Castro-Mondragon et al., 2022), we constructed a TRN governing spike formation after floral transition. The TRN involves numerous TFs, including well-studied factors like VRN1, TaTB1, and TaFUL3 in wheat, as well as TFs from MADS-box, SBP, YABBY, CO-like, AP2-ERF, and NF-YB families known to regulate inflorescence formation in other crops (Yuan et al., 2020; Kellogg, 2022; Luo et al., 2023). Combining publicly available GWAS data with a focus on spike development traits (Hao et al., 2020; Pang et al., 2020; Li et al., 2022), we identified 227 TFs, including 42 that have been functionally analyzed in wheat or other crops. We confirmed the functionality of 36 novel TFs showing potential spike development defects, with further validation of ten TFs through backcross and co-segregation analysis of mutants in these genes. The availability of sequenced mutant populations in multiple genetic backgrounds (Krasileva et al., 2017; Wang et al., 2023) offers an efficient way of screening regulators of agronomic traits in crops. To advance our understanding, we encourage further exploration of TRNs in inflorescence development across different cereals, including barley, oats, and rye, as well as the ’model’ cereal species *Brachypodium distachyon* (Girin et al., 2014). Conducting comparative analyses will help identify novel regulators with excellent alleles present in these underutilized germplasm resources, leading to insights into the varied inflorescence structures, such as the differences in spike or spikelet termination and indeterminacy between barley and wheat (Koppolu and Schnurbusch, 2019). Cases like *DUO-B1* demonstrate the potential for such discoveries (Wang et al., 2022). By unraveling these regulatory networks, we can gain valuable insights for crop improvement and optimization of inflorescence architecture.

### Novel regulator for wheat spike development with excellent allele selected during domestication and breeding process in China

The TRN not only aids in identifying novel factors involved in inflorescence development but also provides valuable guidance for gene functional studies. An exemplary case is the hierarchical regulatory module involving known factors such as TaSPL15-TaAGLG1-TaFUL2. By leveraging the TRN, we also facilitated the functional study of novel factors, identifying their upstream regulators and downstream targets, as demonstrated with TaMYB30-A1. The TRN also enabled us to identify critical regions involved in transcriptional regulation with high resolution. For instance, in the case of *TaMYB30-A1*, we discovered that the excellent haplotype containing a specific SNP in the binding motif of the upstream regulator WFZP was selected during domestication in both tetraploid and hexaploid wheat, as well as in the breeding process in China.

Improving GNPS has been a primary breeding target for enhancing grain yield potential, with different breeding strategies employed in various agro-ecological zones. Under this premise, the C-allele of *TaMYB30-A1*, which exhibits slightly reduced SNS but increased spikelet fertility and higher final GNPS, was selected, resulting in a significant increase in the frequency of this excellent allele. A similar strategy could be applied to pinpoint beneficial alleles of other key regulators for breeding selection, and precision genome editing could be employed to stack multiple excellent loci. Notably, the direction of spike architecture selection may vary for different agro-ecological zones, necessitating testing of cis-element mutations of key factors in different backgrounds to enhance grain yield.

### A valuable database for studying genetic regulation of wheat spike formation

Two recent studies, Wheat-RegNet (Tang et al., 2023) and wGRN (Chen et al., 2023), integrate gene expression, chromatin status and GWAS analysis to predict gene regulation in wheat. While important, the limitations of these studies include the lack of tissue specificity and in-depth validation of their predictions *in planta*. In contrast, our approach involves the development of a gene regulatory network based on RNA-seq and ATAC-seq data from the same tissues at different stages of wheat spike development. We have rigorously validated our predictions through mutant and transgenic plant experiments, as well as in-depth functional assays, thus enhancing the biological relevance of our findings. Our TRN has uncovered novel insights into potential TFs that influence wheat spike architecture and facilitate breeding selection processes.

To facilitate the use of this data source, we have created the Wheat Spike Multi-Omic Database (WSMOD) (http://39.98.48.156:8800/#/) for visualizing gene-centered data such as stage-specific expression, epigenetic landscape, co-expression network, and TRN (**Supplemental Figure 9**). This database also includes links to numerous spike developmental trait-related GWAS analyses and the KN9204 TILLING mutant library with >98% mutation coverage (Wang et al., 2023). This will create further opportunities to explore the molecular insights of inflorescence development beyond TFs. Looking ahead, to further enhance the accuracy and resolution of our TRN and facilitate gene functional studies, we are currently generating single-cell spatially resolved transcriptome datasets. This will allow us to study the heterogeneity of cell populations with spatial information during spike development, providing an even more comprehensive understanding of the regulatory processes involved.

## Methods

### Plant materials, growth condition and sampling

The winter wheat cultivar KN9204 was used in this study. The germinated seeds were treated at 4 °C for 30 days. The seedlings were transplanted into soil and grown in the greenhouse at 22 °C/20 °C day/night, under long day conditions (16 h light/8 h dark). The stage-specific shoots of wheat were dissected under the stereomicroscope based on the anatomic and morphological features, and immediately frozen in liquid nitrogen and stored at -80 °C. About 10 to 50 spikes were pooled for each biological replicate for RNA-seq (three replicates), ATAC-seq and CUT&Tag (two replicates) analysis at eight or five development stages.

### Generation, identification and phenotyping of transgenic lines and mutants

To obtain RNAi transgenic wheat plants, the specific fragment of *TaSPL15* was separately amplified and inserted into pc336 (*Ubi:GWRNAi:NOS*) vector using Gateway cloning method. The constructed vector was transformed into callus to generate the transgenic plants as described previously (Liu et al., 2023). Primers for genotyping are listed in **Supplemental Table 12**.

Loss-of-function mutations for key TFs were screened in the database of sequenced ethyl methane sulphonate (EMS) mutagenized populations of the hexaploid winter wheat cv. Kenong9204 (Wang et al., 2023) and the Spring wheat cv. Cadenza (Krasileva et al., 2017), which were confirmed using Sanger sequencing (Subgenome-specific amplification primers were described in **Supplemental Table 12**). At least two independent mutant lines in KN9204 background (or both KN9204 and Cadenza background) were selected for each gene, and all these mutants were backcrossed once or twice to wild-type plants to reduce background mutation load prior to phenotyping. The RNAi knockdown transgenic plants of *TaSPL15* and isogenic sibling lines from the backcross population of ten novel TFs were grown at the Experimental Station of Institute of Genetics and Developmental Biology, Chinese Academy of Sciences, Beijing. The heading date was defined as the days from sowing to 50% of the spikes fully emerged from the boot. Spike-related traits were measured 15 days after flowering with 15∼25 prominent tiller spikes. Spike length was measured from the base of the spike to the tip (excluding the awns), and the fertile and sterile spikelet number per spike was counted separately to obtain the SNS, fertile spikelets per spike (FSPS), and SSPS. The spikelet fertility ratio is defined as the ratio of fertile and total spikelet numbers. After maturity, these spikes were harvested and used to measure the GNPS.

### RNA extraction, sequencing, RT-qPCR and *in situ* hybridization

Total RNA was extracted using HiPure Plant RNA Mini Kit according to the manufacturer’s instructions (Magen, R4111-02). RNA-seq libraries’ construction and sequencing platform were the same as previous description (Zhao et al., 2023), by Annoroad Gene Technology.

First-strand cDNA was synthesized from 2 μg of DNase I-treated total RNA using the TransScript First Strand cDNA Synthesis SuperMix Kit (TransGen, AT301-02). RT-qPCR was performed using the ChamQ Universal SYBR qPCR Master Mix (Vazyme, Q711-02) by QuantStudio5 (Applied biosystems). The expression of interested genes was normalized to Tubulin for calibration, and the relative expression level is calculated via the 2^-ΔΔCT^ analysis method (Livak and Schmittgen, 2001). Primers used for RT-qPCR are listed in **Supplemental Table 12**.

RNA *in situ* hybridization was carried out as described previously (Cui et al., 2010). Fresh young spikes were fixed in formalin-acetic acid-alcohol overnight at 4 °C, dehydrated through a standard ethanol series, embedded in Paraplast Plus tissue-embedding medium (Sigma-Aldrich, P3683), and sectioned at 8 μm width using a microtome (Leica Microsystems, RM2235). Digoxigenin-labeled RNA probes were synthesized using a DIG northern Starter Kit (Roche, 11277073910), according to the manufacturer’s instructions. Primers used for probe synthesis are listed in **Supplemental Table 12**.

### Data Preprocessing and reads alignment

Raw reads were filtered by fastp v0.20.1 with parameter “--detect_adapter_for_pe” for adapters removing, low-quality bases trimming, and reads filtering (Chen et al., 2018). Furthermore, FastQC v0.11.8 (http://www.bioinformatics.babraham.ac.uk/projects/fastqc/) was performed to ensure the high quality of reads.

Reads were aligned using either BWA-MEM v0.7.17 with parameter “-M” (for ATAC-seq and CUT&Tag seq) or hisat2 v2.1.0 with default parameters (for RNA-seq) to the reference genome of *Triticum aestivum* cv. Chinese Spring (IWGSC RefSeq v1.0, https://urgi.versailles.inra.fr/download/iwgsc/IWGSC_RefSeq_Assemblies/v1.0/) (Appels et al., 2018; Kim et al., 2019; Li and Durbin, 2009). Gene models from the IWGSC Annotation v1.1 were used as the reference and high-confidence genes were used throughout this study. The resulting SAM files were converted to BAM format, sorted, and indexed using Samtools v1.4 (Danecek et al., 2021). Sam files of RNA-seq generated from hisat2 were converted to bam files without deduplication. For ATAC-seq and CUT&Tag, SAM files were further filtered with “samtools view -bS -F 1,804-f 2 -q 30” to filter out the low-quality mapped reads. Duplicates in the high-quality mapped reads were removed using Picard v2.23.3. Two replicates bam files were merged by samtools. To normalize and visualize the individual and merged replicate datasets, the BAM files were converted to bigwig files using bamCoverage provided by deepTools v3.3.0 with 10 bp bin size and normalized by RPKM (Reads Per Kilobase per Million mapped reads) with parameters “-bs 10 --effectiveGenomeSize 14,600,000,000 --normalizeUsing RPKM --smoothLength 50” (Ramirez et al., 2014).

### RNA-seq data analyses

The number of paired reads that mapped to each gene was counted using feature Counts v2.0.1 with the parameter "-p" (Liao et al., 2014). The counts files were then used as inputs for DEGs (differentially expressed genes) analysis by DESeq2 v1.26.0 with a threshold "|Log2 Fold Change| ≥ 1 and FDR ≤ 0.05" (Love et al., 2014). The raw counts were further normalized to TPM (Transcripts Per Kilobase Million) for gene expression quantification. For subsequent clustering and visualization, we obtained mean counts by merging three biological replicates. TPM values of genes were Z-scaled and clustered by k-means method and displayed using R package ComplexHeatmap (v2.4.3) (Gu et al., 2016). Functional enrichment was performed using an R package clusterProfiler v3.18.1, and GO annotation files were generated from IWGSC Annotation v1.1 (Yu et al., 2012).

### Cut&Tag and ATAC-seq experiment and data analyses

ATAC-seq and CUT&Tag experiments followed the previously described method (Zhao et al., 2023). Tn5 transposase used and tagmentation assay is done following the manual (Vazyme, TD501-01). Libraries were purified with AMPure beads (Beckman, A63881) and sequenced using the Illumina Novaseq platform at Annoroad Gene Technology. Antibodies used for histone modifications are listed in **Supplemental Table 13**.

Data processing and reads alignment were performed as previously described (Zhao et al., 2023). MACS2 v2.1.4 was used to call peaks. Parameters “-p 1e-3” was used for H3K27ac, H3K4me3 and H2A.Z; parameters “--broad –broad-cutoff 0.05” were used for H3K27me3 and H3K36me3 (Zhang et al., 2008). For ATAC-seq data, MACS2 was used with parameters “--cutoff-analysis --nomodel --shift -100 --extsize 200”. The peaks were annotated by R package ChIPseeker v1.26.2 with "annotatePeak" function (Yu et al., 2015). The gene promoters are defined as 3.0 Kb upstream of gene TSS. For the identification of transcription factor footprints in ATAC-seq peaks, we used the HINT tool v0.13.2 of the Regulatory Genomics Toolbox (RGT) (Gusmao et al., 2014). The custom wheat genome was configured using IWGSC refseq v1.1 Chinese Spring genome based on the introduction of HINT software. TF motifs were downloaded from the JASPAR Plantae database (https://jaspar.genereg.net/) (Castro-Mondragon et al., 2022).

### Differential chromatin modification enriched regions detection

For Cut&Tag and ATAC-seq, reads count and CPM normalized value of peaks were calculated by R package DiffBind v2.16.2 with the setting "DBA_SCORE_TMM_READS_EFFECTIVE_CPM". DiffBind was also used to identify differentially accessible regions (DARs) and histone modification enriched regions with parameters "method = DBA_DESEQ2" and a threshold "|Log2 Fold Change| ≥ 1 and FDR ≤ 0.05".

### Pseudotime indexing and gene regulatory network construction

We used the pseudotime indexing method to analyze gene expression as described in previous studies with some modifications (Hao et al., 2021; Leiboff and Hake, 2019; Zhao et al., 2023). All of the expressed genes during spike reproductive development (Stages from W3 to W4) were used to separate samples on a PCA plot. Then, each developmental stage was assigned a location and the Euclidean distance between adjacent stages was calculated and scaled from 0.0 to 10.0. For each gene, we calculated the fitted curve and interpolated the curve into 500 points based on gene expression using the “loess” function in R. We further performed PCA for each gene based on the standardized expression data and used “atan2” function in R to order genes based on the time of expression.

For TRNs construction, we only focused on DEGs with TPM ≥ 0.5 in at least one stage from W3 to W4. The potential upstream regulatory TFs of one gene were predicted based on the motif present at its regulatory region. Here, HINT tool v0.13.2 were used to identify footprints within ATAC-seq peaks and motifs within the footprints, and then matched the motifs to TFs based on JASPAR Plantae database. The TFs in wheat identified by blastp (v 2.10.1) (Camacho et al., 2009) with criteria “e value < 1e-10 and identity > 40%” using TFs of JASPAR Plantae database (Castro-Mondragon et al., 2022).The regulatory relationships were further filtered following criteria of “the TF and target gene were co-expressed (supported by WGCNA network)

And both with TPM ≥ 0.5 at least one stage from W3 to W4” to obtain the TF-target regulation. We used *k*-means function in R to cluster genes into six categories and performed hypergeometric test to calculate *P*-value of regulation among gene categories.

### Genome-wide and Gene-based association analysis

A previously characterized population (Li et al., 2022), consisting of the Chinese wheat mini-core collection (MCC, 262 accessions) and 25 currently widely grown cultivars, was used to investigate the genetic basis underlying SSPS variation in wheat. Field experiments were carried out in Beijing and Shijiazhuang (Hebei province, China) during three successive cropping seasons (2019-2022), using a completely randomized design with two repetitions. Sixteen seeds of each accession were planted in a 150-cm-long row with 23 cm between rows. At maturity, spikes of the main stem from ten plants in the center of the row were used for investigates the spike characters. Spikelet without any grains was counted as sterile ones, and spikelet produce grains regarded as fertile ones. The average number of fertile spikelets and sterile spikelets for ten spikes were defined as the FSPS and SSPS, respectively. The spikelet fertility ratio was defined as “FSPS / (FSPS + SSPS)”. The best linear unbiased prediction (BLUP) value for the SSPS trait was performed in GWAS analysis, with 984,034 high quality SNPs (missing rates ≤ 0.2, MAF ≥ 0.05) captured by exome sequencing (Li et al., 2022), using the EMMAX software package (Kang et al., 2010). The *P* value of 1E-04 was determined to be the suggestive threshold for declaring a significant association according to the adjusted Bonferroni method.

All nucleotide polymorphisms (unfiltered SNP and short InDels) in 6 Kb genomic region of *TaMYB30-A1*, including exons, intron regions, 4 Kb promoter regions and 0.5 Kb 3’-UTR regions, were performed to the gene-based association with SSPS using Tassel v5.2. Haploview 4.2 was used to calculate the pairwise linkage disequilibrium coefficient (*r^2^*) between SNPs and draw the LD plot. Manhattan plots and quantile-quantile plots were generated using R package “CMplot” (https://github.com/YinLiLin/R-CMplot).

### Spike morphology observation by scanning electron microscopy (SEM)

Photomicrographs of young spikes were taken using a stereomicroscope (Leica Microsystems, S8 APO) equipped with a digital camera (Canon, A640). For SEM, young spikes from each stage were fixed in 2.5% glutaraldehyde at 4°C. After dehydration in a series of ethanol solutions and substitution with 3-methylbutyl acetate, the samples were subjected to critical point drying, coated with platinum, and observed using a variable pressure scanning electron microscope (Hitachi S-3000N).

### Luciferase (LUC) reporter assay

For LUC analyses, full-length coding sequences of *TaSPL15*, *TaAGLG1*, *TaFUL2*, *TaMYB30-A1*, *WFZP-D1* and *TaDOF17-B1* were cloned into PTF101 vector to generate the effector construct *35S: TF*-GFP; About 3 Kb promoter fragment of *TaAGLG1*, *TaFUL2*, *TaHMA*, *TaMYB30-A1*, *TaAP2-39-D1*, *TaBAK1-D1* and *TaPILS7-A1* were amplified and fused in-frame with the CP461-LUC vector to generate the reporter construct *proGOI*: LUC (see **Supplemental Table 12** for primers), and point mutation at TF binding motifs were induced in *proGOI:LUC* to generate the *proGOIm:LUC*. The plasmids were transformed into *Agrobacterium* GV3101. The mixture of bacterial solution *35S*:TF-GFP or *35S*: GFP (OD = 0.5), *proGOI:LUC* or *proGOIm:LUC* (OD = 0.5) and P19 (OD = 0.3) in activation buffer (10 mM MES, 150 μM AS, 10 mM MgCl_2_) was injected to tobacco (*Nicotiana benthamiana*) leaf. pSUPER-GFP, target-pro:LUC and P19 as control. Firefly luciferase (LUC) and Renilla luciferase (REN) activities were measured using dual luciferase assay reagent (Promega, VPE1910) after one day ’co-cultivation in dark and two days in light, the relative value of LUC/REN is indicated as average with standard error of multiple replicates.

### The establishment of Wheat Spike Multi-Omics Database (WSMOD)

The Wheat Spike Multi-Omics Database (http://39.98.48.156:8800/<u>#/</u>) integrates a core transcription regulatory network during wheat spike development, encompassing multi-omics data such as gene expression, chromatin accessibility, multiple histone modifications, SNP information, and spike-related GWAS signals (**Supplemental Figure 8**). Additionally, a co-expression network has been constructed using the expression data of eight stages during spike development. Users can access the expression, regulatory network, and other relevant information by inputting the gene ID via six functional modules: Search by Gene ID, WGCNA, TRN, Epigenomic Status, GWAS, and KN9204 EMS Mutants. For search by gene ID function, WSMOD provides gene expression patterns, orthologs in rice and *Arabidopsis thaliana*, open chromatin regions and footprint, and other information. WGCNA function provides gene-gene co-expression relationships and network visualization. TRN function provides information of genes upstream TFs, TF binding sites and motifs, as well as downstream target genes. Epigenomic status function offers genome visualization of transcription, chromatin accessibility, various histone modifications, variations and QTL sites. GWAS function provides spike-related QTL regions and gene information in regions. KN9204 EMS Mutants function provides a quick link to KN9204 TILLING database in which users can conveniently query and order the EMS mutants of the interested gene. All search results are presented in images and interactive tables that can be downloaded for further analysis. These multi-omics data sources and transcription regulatory networks would deepen our understanding of genetic regulation in wheat spike development and provide information for gene function analysis.

### Statistics and data visualization

R (https://cran.r-project.org/;version 4.0.2) was used to compute statistics and generate plots if not specified. The Integrative Genomics Viewer (IGV) was used for the visual exploration of genomic data (Robinson et al., 2011). The correlation between two groups of data was conducted with the Pearson analysis (**Figures 2B, 2E, 7I and Supplemental Figure 8E**) and *P* values are determined by the two-sided Pearson correlation coefficient analysis (**Figure 2E**). For enrichment analysis, Hypergeometric test (**Figures 2C, 3D**) and Fisher’s exact test were used (**Figures 2F, 3C and Supplemental Figures 3D, 3E**). For two groups ’comparison of data that fit a normal distribution, the Student’s two-tailed t-test was used (**Figures 3G, 3H, 5E, 6C, 6D, 6F, 6G, 7E, 7F and Supplemental Figures 4G, 5B, 5C, 5D, 7A,7B, 8D**). For two groups ’ comparison of data that does not fit a normal distribution, the Wilcoxon rank-sum test was used (**Figures 4A, 4F, 7C,7G,7J and Supplemental Figures 6A, 8B, 8F**).

## Data availability

The raw sequence data reported in this paper have been deposited in the Genome Sequence Archive (Chen et al., 2021) in National Genomics Data Center (CNCB-NGDC Members and Partners, 2022), China National Center for Bioinformation / Beijing Institute of Genomics, Chinese Academy of Sciences (CRA008877) that are publicly accessible at https://ngdc.cncb.ac.cn/gsa.

## Code availability

Code used for all processing and analysis is available at GitHub (https://github.com/yx-xu/wheat-spike-development).

## Funding

This research is supported by the National Natural Sciences Foundation of China (31921005), Strategic Priority Research Program of the Chinese Academy of Sciences (XDA24010204), National Key Research and Development Program of China (2021YFD1201500), and the Major Basic Research Program of Shandong Natural Science Foundation (ZR2019ZD15).

## Author contributions

J.X. designed and supervised the research, J.X., X.-L. L., Y.-X.X., D.-Z.W. wrote the manuscript. X.-L. L. did the sample collection and *in situ* hybridization; X.-L. L. and X.-Y.Z. did plasmid construction, RT-qPCR and co-segregation experiment; X.-M. B. did wheat transformation; X.-L. L. and Y.-M.Y. performed CUT&Tag, ATAC-seq and RNA-seq experiments; X.-Y. Z. did the reporter assay; Z.-X. C. generated the website of WSMOC for visualization; Y.-L. D., L.M., X.-Y. Z., F.L., X.-S. Z., C.U. and X.-D. F. provide some raw data or plant materials; Y.-X. X., D.-Z. W. performed bio-informatics analysis; C.U. and X.-D. F. revised the manuscript; X.-L. L., Y.-X. X., D.- Z. W., Y.-M. Y. and J.X. prepared all the figures. All authors discussed the results and commented on the manuscript.

## Supporting information

Supplemental Table 1. The expression and k-means clusters of genes

Supplemental Table 2. Number of DEGs between samples

Supplemental Table 3. DAPR identified from SAM to W3

Supplemental Table 4. Chromatin accessibility and k-means clusters of DAPR from SAM to W3

Supplemental Table 5. Gene list of three gene sets

Supplemental Table 6. The expression and k-means clusters of DEGs from W3 to W4

Supplemental Table 7. Reported spike-related QTLs used in this study

Supplemental Table 8. Identified key factors involved in spike development

Supplemental Table 9. Classification and mutant lines of 227 candidate genes

Supplemental Table 10. Summary of the backcross for identified ten novel TFs

Supplemental Table 11. SNPs significantly associated with SSPS in GWAS

Supplemental Table 12. List of researched genes related to wheat spike development

Supplemental Table 13. Primers used in this study

Supplemental Table 14. Antibodies used in this study

## Acknowledgements

We thank Dr. Y.L. Jiao (College of Life Sciences, Peking University) for the *TaAGLG1*-OE transgenic wheat materials, Dr. X.G. Liu (College of Life Sciences, Hebei Normal University) for the *WFZP*-OE transgenic wheat materials, and Drs. Z.F. Ni and J. Liu (College of Agronomy and Biotechnology, China Agricultural University) for the *WFZP* CRISPR mutants.

## Competing interests

The authors declare no competing interests.

## Supplemental Figures

**Supplemental Figure 1.**
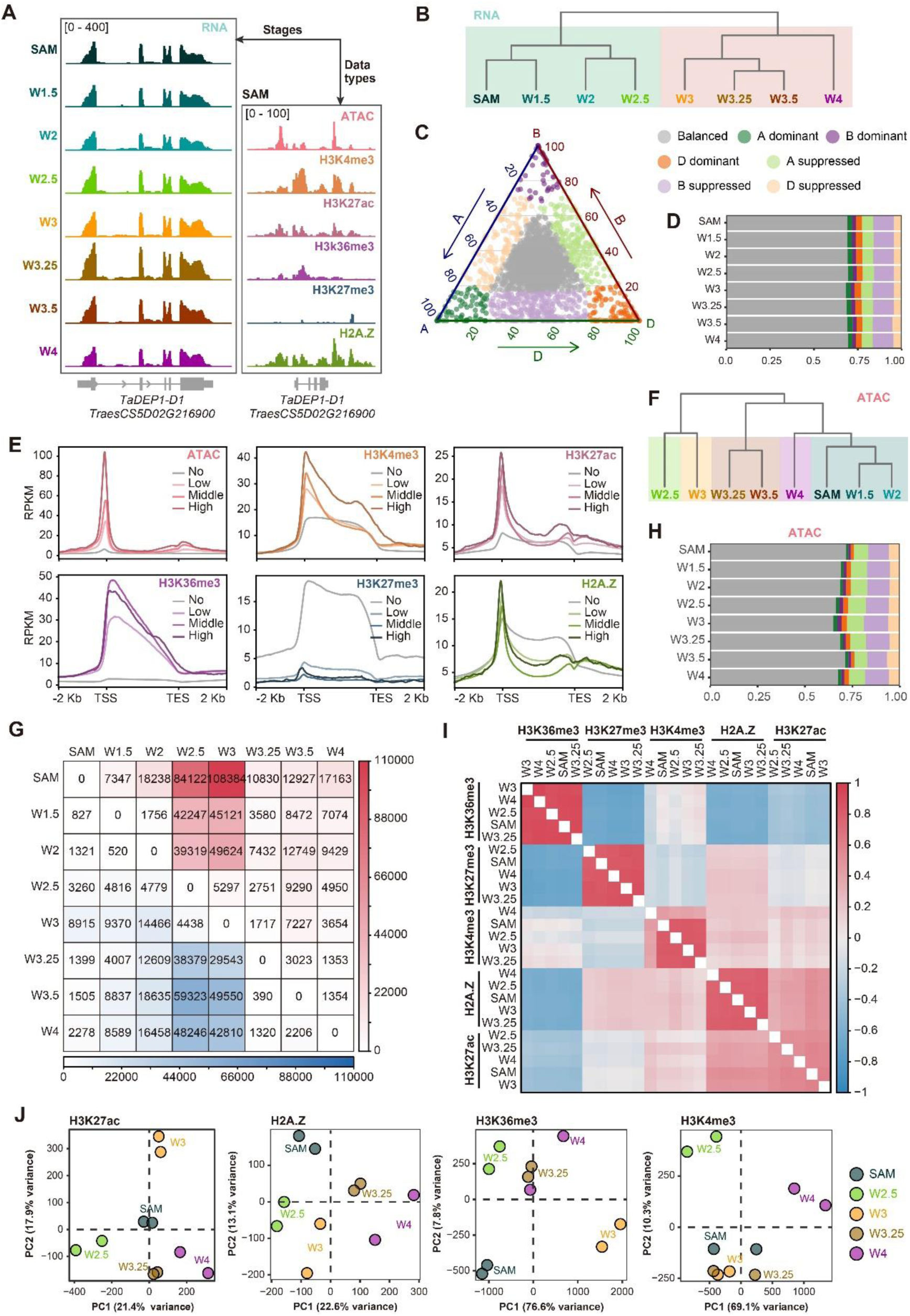
Features of histone modifications during spike development. **A,** Example browser view at *TaDEP1* locus showing epigenomic data types at SAM stage and transcriptome levels of different sampling stages. RPKM was used for data normalization. **B,** Cluster dendrogram of RNA-seq data showing two distinct development clusters: vegetative to flowering transition cluster (SAM, W1.5, W2 and W2.5 stages) and reproductive growth phase cluster (W3, W3.25, W3.5 and W4 stages). **C,** Ternary plot showing homoeologs expression bias of triads in hexaploid wheat. Each circle represents a gene triad with an A, B, and D coordinate consisting of the relative contribution of each homoeolog to the overall triad expression. The colors of the circle indicate different expression bias types. **D,** Proportion of triads in each category of homoeolog expression bias across the spike developmental stages. **E**, Metagene profile for histone modifications of gene sets with different expression levels. No, Low, Middle and High represent gene expression levels. **F,** Cluster dendrogram of ATAC-seq data showing five distinct development clusters: vegetative cluster (SAM, W1.5, W2 stages), flowering transition stage (W2.5 stage), inflorescence initiation (W3 stage), spikelet formation (lemma primordium, W3.25 and floret primordium visible, W3.5), terminal spikelet formation (terminal spikelet and stamen primordium formation, W4 stage). **G,** The matrix of differentially accessible regions (DARs) across developmental stages. The number of decreased and increased chromatin accessibility compared with previous stages was represented in the lower-triangle (number in light blue) and upper-triangle panel (number in light red), respectively. A region with |log_2_(Fold Change)| ≥ 1 and FDR ≤ 0.05 by DiffBind between any two stages was considered as DAR. **H,** Proportion of triads in each category of homoeolog chromatin accessibility (gene promoter region) bias across the spike developmental stages. **I,** Pair-wise correlation map among different histone modification profiles. Jaccard index was calculated based on the peaks overlap, and then Pearson correlation scores were generated. **J,** PCA of H3K27ac, H2A.Z, H3K36me3 and H3K4me3 samples during spike development. Each dot represents one sample; two biological replicates are sequenced for each stage.

**Supplemental Figure 2.**
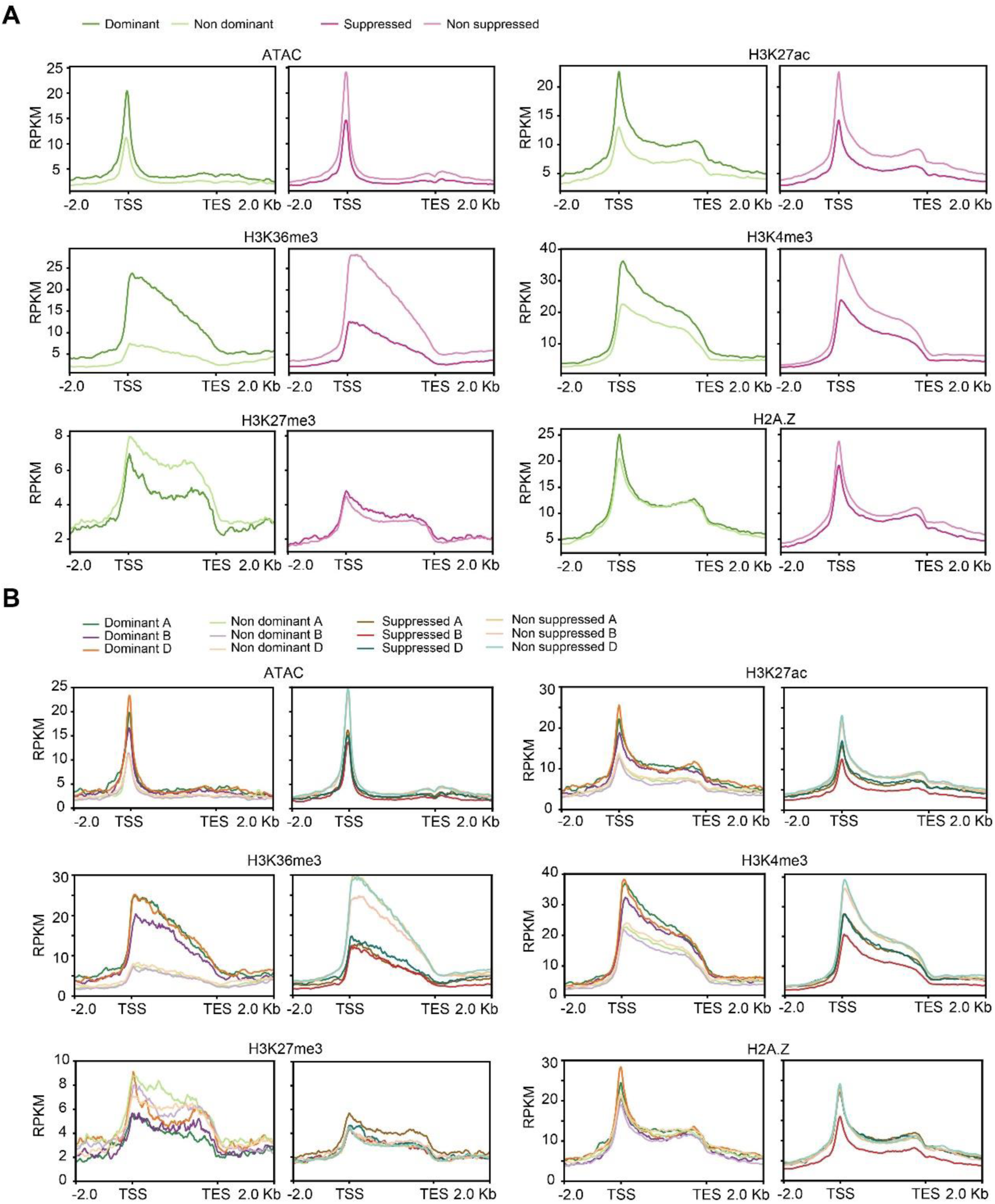
Chromatin landscape of unbalanced expressed genes. **A**, Metagene profile for chromatin accessibility and histone modifications around the gene body for unbalanced triads. The dominant triads were separated into dominant (green) and non dominant (pale green) homoeologs, and suppressed triads separated into suppressed (purple) and non suppressed (pale purple) homoeologs. Epigenetic modification signals were normalized using RPKM, with 10 bp bin size. **B**, Metagene profile for chromatin accessibility and histone modifications around the gene body for subgenome dominant, non dominant, suppressed, and non suppressed triads genes. Dominant and suppressed homoeologs of A, B, and D subgenomes were represented in green; purple; and orange, respectively. Non-dominant and non-suppressed homoeologs of A, B, and D subgenomes were represented in pale green; pale purple; and pale orange, respectively.

**Supplemental Figure 3.**
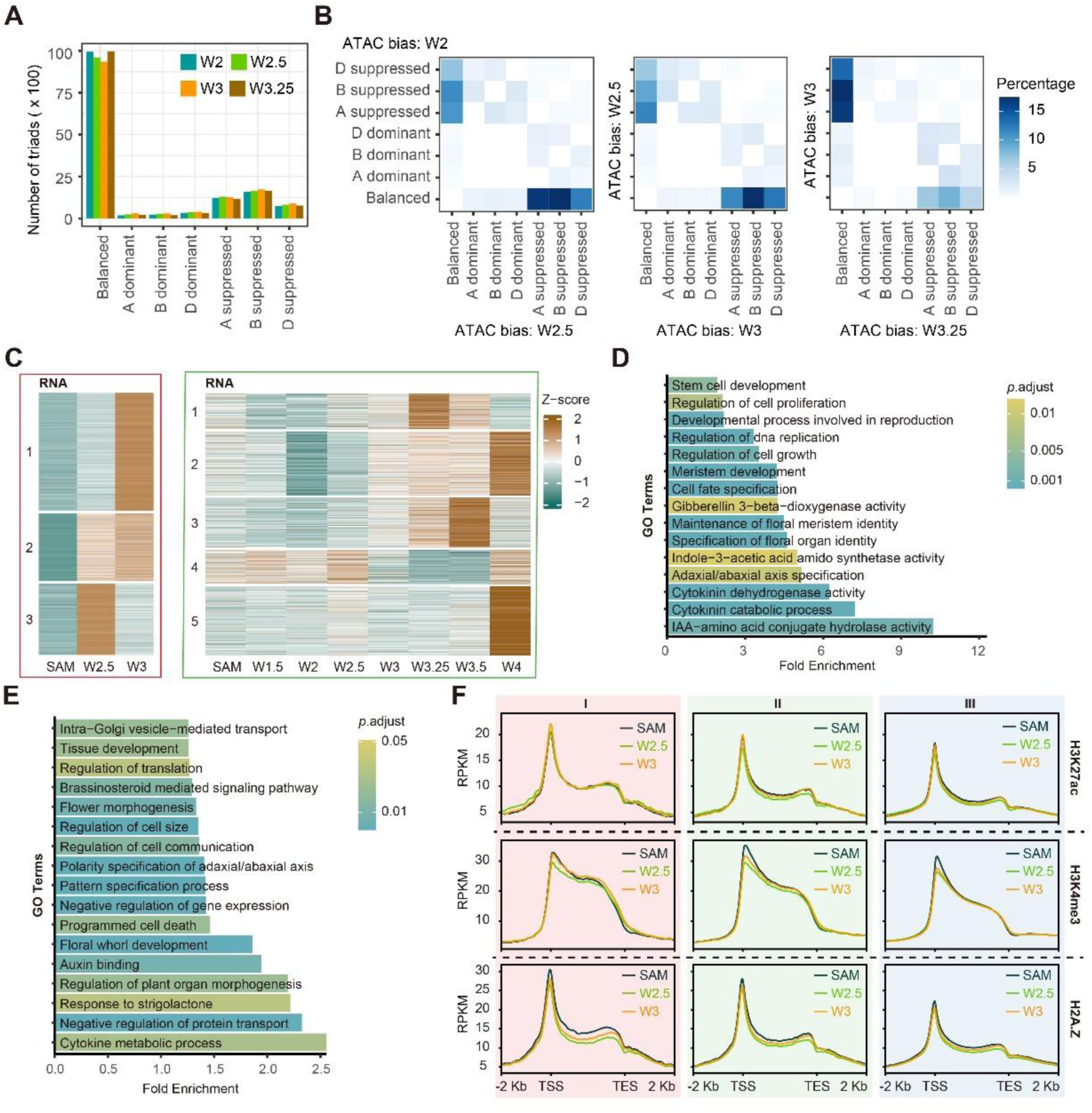
Chromatin landscape dynamics associated with transcriptional change from vegetative-to-reproductive transition. **A,** Number of triads in each chromatin accessibility bias group at four stages. W2, W2.5, W3, and W3.25 stages were marked in teal, green, orange, and brown, respectively. **B,** Percentage of triads whose chromatin accessibility bias changed from W2 to W2.5, from W2.5 to W3, and from W3 to W3.25. Triads in which chromatin accessibility bias did not change were not counted. **C,** Expression pattern of genes up-regulated at W2.5 and W3 stages versus SAM stage (left, red frame) and genes up-regulated at later stages rather than W2.5 or W3 stage (right, green frame). Heatmap showing *k*-mean clustering of gene expression. **D-E,** GO enrichment analysis of genes in gene set Ⅱ (**D**), Ⅲ (**E**). **F,** H3K27ac (top), H3K4me3 (middle) and H2A.Z (bottom) levels of genes in gene set Ⅰ, Ⅱ, Ⅲ at SAM, W2.5 and W3 stages. The Y axis indicates RPKM normalized values of read density.

**Supplemental Figure 4.**
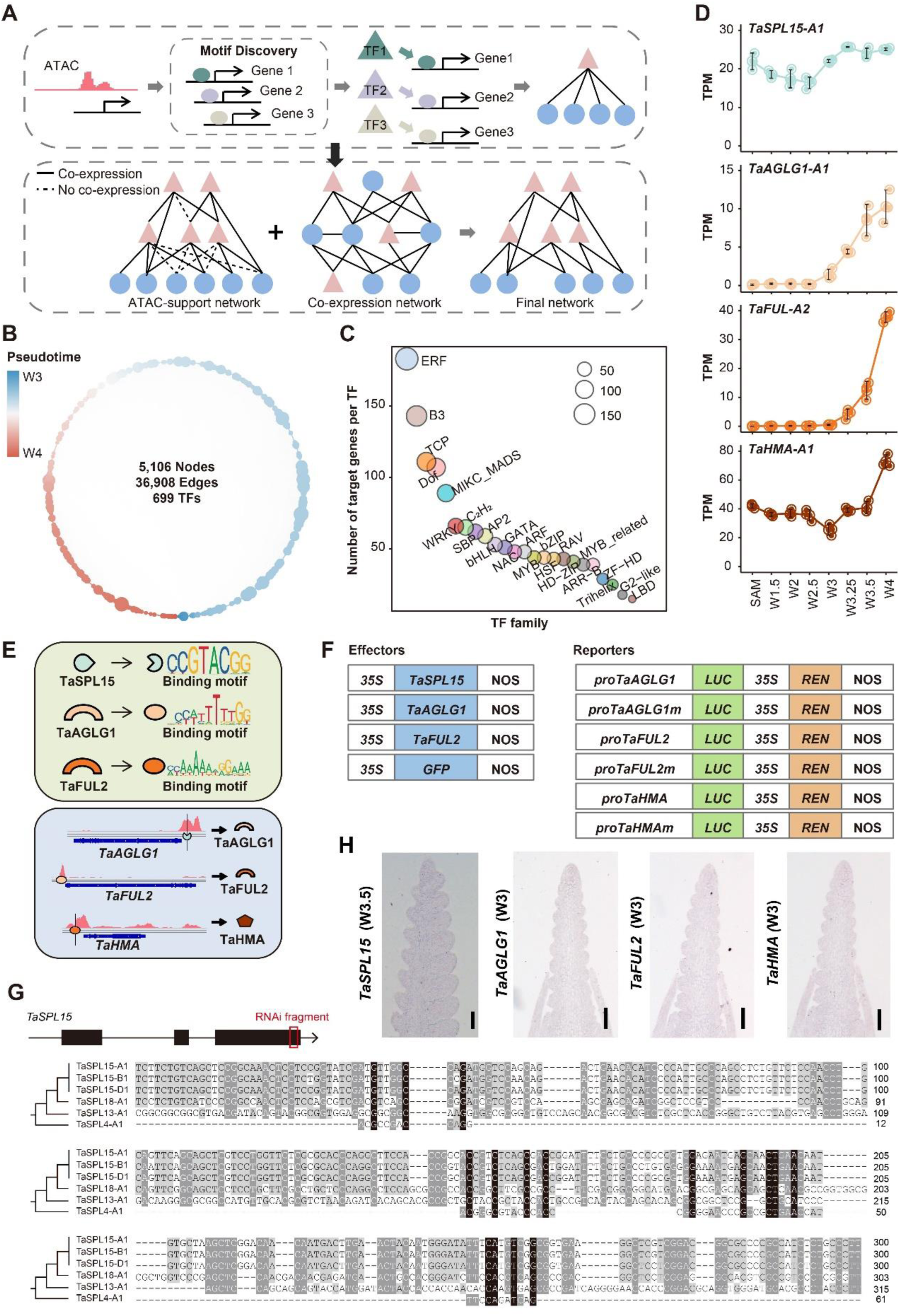
Construction of transcription regulatory network (TRN). **A,** Schematic of the strategy for TRNs building (see Methods for detail). **B,** Total TRNs characterized for governing spike architecture formation. **C,** Average number of target genes of each TF family in the TRN. The circle size represents number of target genes. **D,** Dynamic expression profile of *TaSPL15-A1*, *TaAGLG1-A1*, *TaFUL-A2* and *TaHMA-A1* at different stages during spike development. **E,** Presence of different TF binding motifs in the target gene’s open chromatin regions from the TaSPL15-TaAGLG1-TaFUL2-TaHMA regulatory module. **F,** Schematic diagram showing the vectors used in the Luciferase reporter assays of Figure 3G. **G**, Schematic diagram and the sequence showing the selection of RNAi interference fragment for specific targeting *TaSPL15*. **H**, Sense probes used in **Figure 3F**. Scale bars = 100 μm.

**Supplemental Figure 5.**
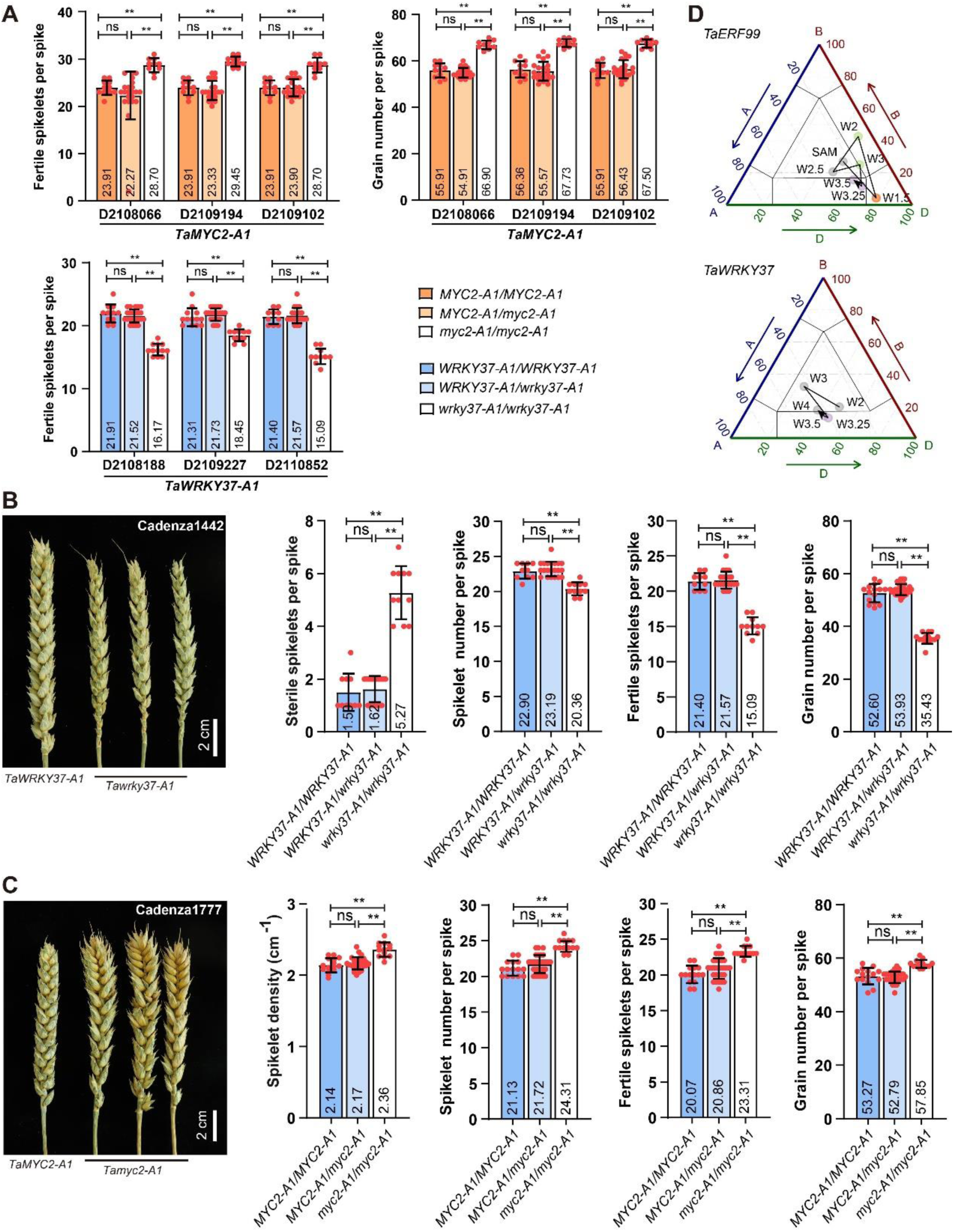
The quantification data of spike phenotype associated with novel factors mutations. **A**, Quantification of FSPS and GNPS for *Tamyc2-A1*, and FSPS for *Tawrky37-A1* in KN9204 TILLING lines. The sibling homozygous mutant plants, heterozygous plants, and wild-type plants identified in the backcross population were analyzed for their phenotype. The error bars denote ±SD, and individual plant data is represented by a dot. Student’s *t*-test was used for the statistical significance. *, *P* ≤ 0.05; **, *P* ≤ 0.01; ns, no significant difference. **B**, The spike developmental defect and quantification of SSPS, SNS, FSPS and GNPS for *Tawrky37-A1* in Cadenza mutant (Cadenza1442) backcross sibling lines. The error bars denote ±SD, and data of each plant is presented. Student’s *t*-test was used for the statistical significance. *, *P* ≤ 0.05; **, *P* ≤ 0.01; ns, no significant difference. **C**, The spike developmental defect and quantification of SD, SNS, FSPS and GNPS for *Tamyc2-A1* in Cadenza mutant (Cadenza1777) backcross sibling lines. The error bars denote ± SD, and data of each plant is presented. Student’s *t*-test was used for the statistical significance. *, *P* ≤ 0.05; **, *P* ≤ 0.01; ns, no significant difference. **D,** Ternary plots showing expression bias of *TaERF99* triad (upper panel) and *TaWRKF37* triad (bottom panel) during spike development. The black arrows show the sequence of development stages of the wheat spike.

**Supplemental Figure 6.**
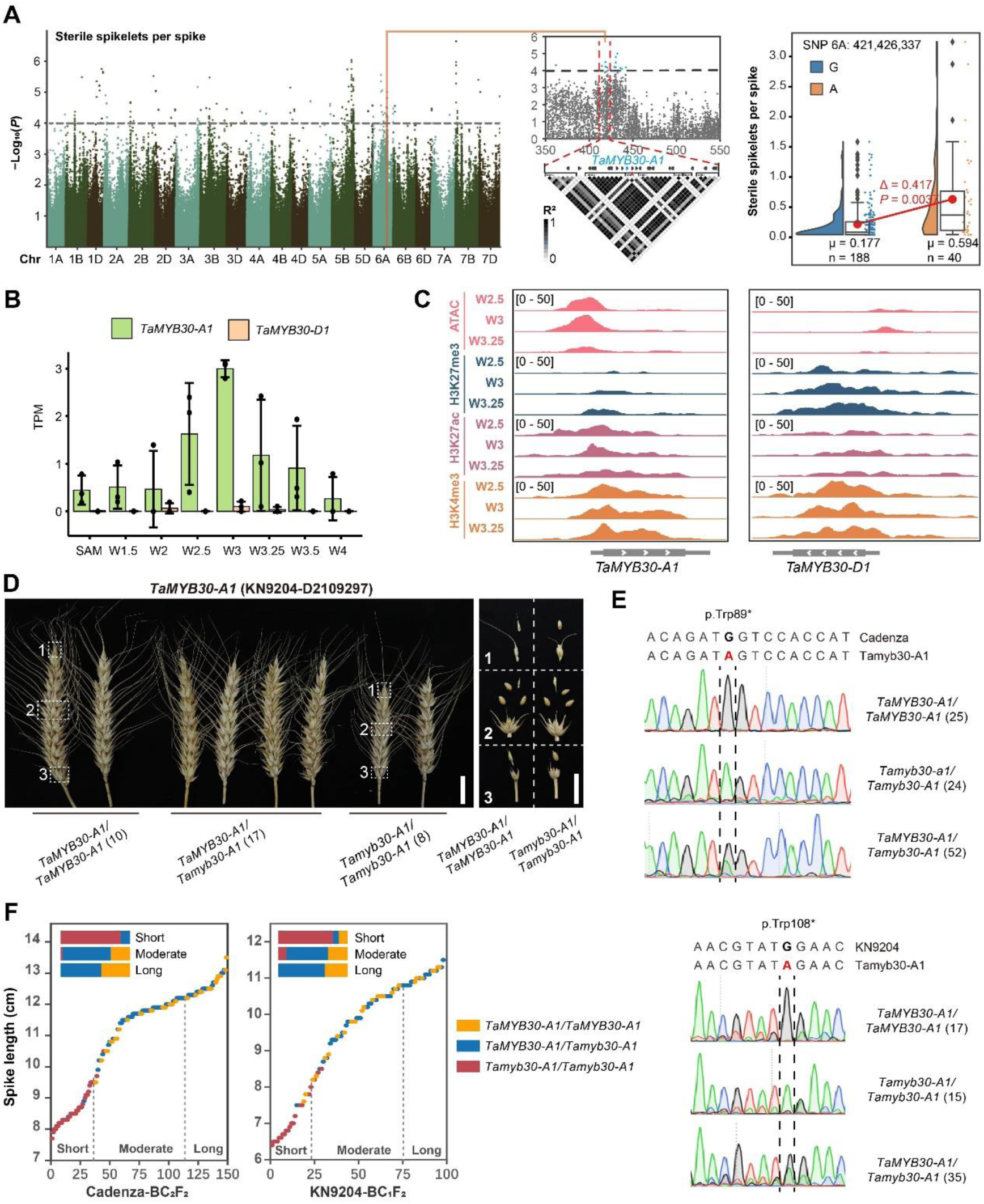
The identification of *TaMYB30-A1* mutants. **A,** Manhattan plot showing the significant GWAS signals for SSPS on chromosome 6A. Dashed horizontal line indicates the genome-wide significance threshold (-log_10_ *P* = 4.0). Local Manhattan plot of SNPs in chr6A: 350-550 Mb is shown, with a LD heatmap showing the LD of 6 Mb physical interval flanking the peak SNP 6A:421426337. The distribution and comparison of SSPS between haplotypes defined by the peak SNP was shown in a raincloud plot. The bars within raincloud box plots represent 25^th^ percentiles, medians, and 75^th^ percentiles. The mean values of the two haplotypes are linked by a red line, and Wilcoxon rank-sum test was used to determine the significance of SSPS between two haplotypes. **B**, The expression level of *TaMYB30* homoeologs across developmental stages. The error bars denote ±SD, and data of each replicate is presented. **C**, Genomic tracks showing expression, chromatin accessibility and histone modifications change at representative genes of *TaMYB30-A1* and *TaMYB30-D1*. **D**, The spike photograph showing the SL, SNS, and spikelet fertility at each part of spike in KN9204 backcross sibling lines of *Tamyb30-A1*. **E**, Mutant genotypes of *Tamyb30-A1* (Cadenza mutant and KN9204 mutant) were confirmed by Sanger sequencing, and the ratio was consistent with wild-type: heterozygote: homozygous = 1:2:1. **F**, The homozygous *Tamyb30-A1* mutant plants were distributed in the smallest quartile of spike length in both BC_2_F_2_ of Cadenza and BC_1_F_2_ of KN9204.

**Supplemental Figure 7.**
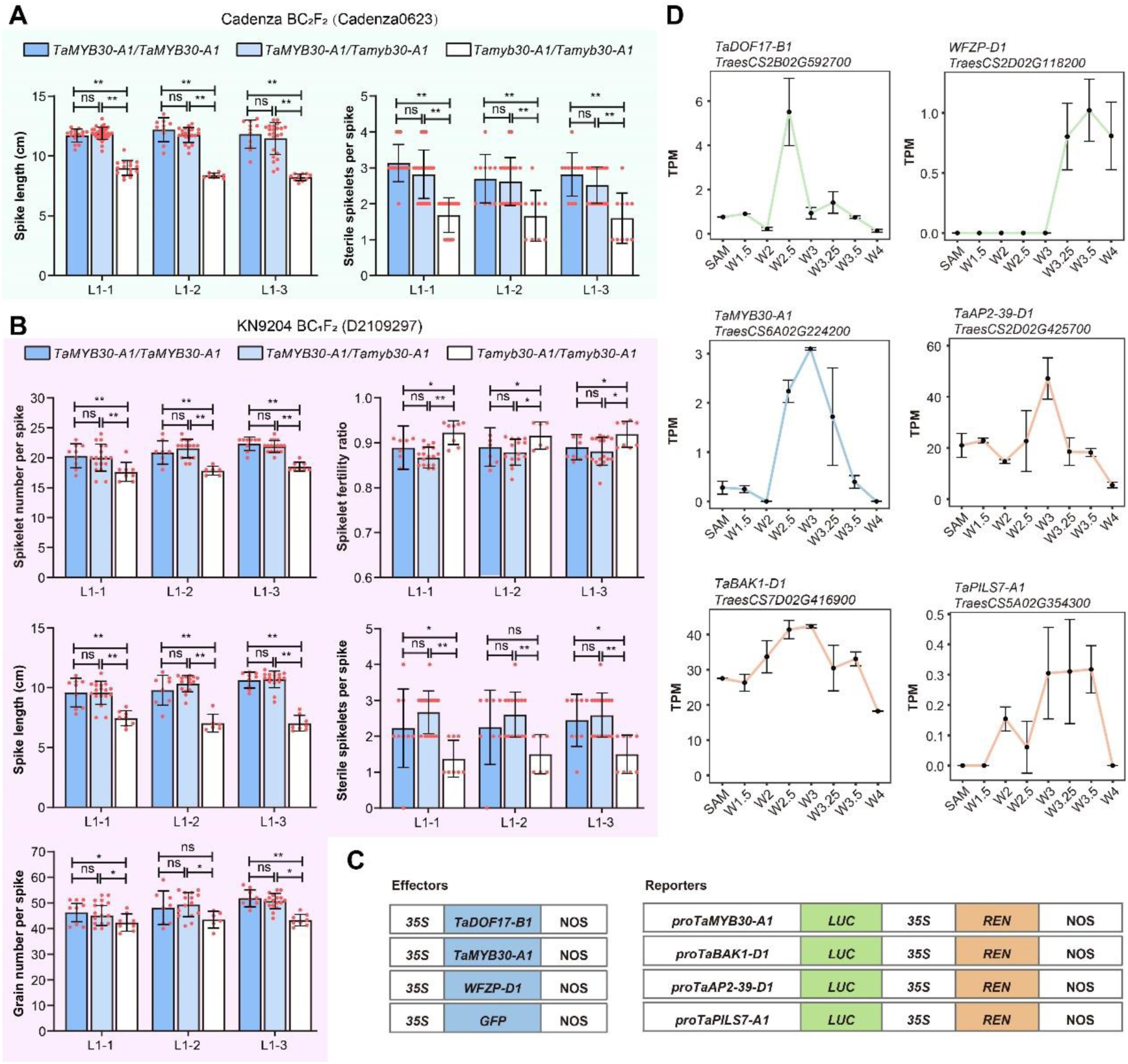
The characterization of *TaMYB30-A* mutants. **A**, Statistical comparison of SL and SSPS for *Tamyb30-A1* in Cadenza backcross sibling lines. L1-1, L1-2 and L1-3 represent independent sibling plants from the same mutant line Cadenza0623. The error bars denote ± SD, and data of each plant is represented by a dot. Student’s *t*-test was used for the statistical significance. *, *P* ≤ 0.05; **, *P* ≤ 0.01; ns, no significant difference. **B**, Statistics comparison of SNS, spikelet fertility, SL, SSPS and SSPS for *Tamyb30- A1* in KN9204 backcross sibling lines. L1-1, L1-2 and L1-3 represent independent sibling plants from the same mutant line D2109297. The error bars denote ±SD, and data of each plant is represented by a dot. Student’s *t*-test was used for the statistical significance. *, *P* ≤ 0.05; **, *P* ≤ 0.01; ns, no significant difference. **C**, Schematic diagram showing the vectors used in the luciferase reporter assays of TaMYB30-A regulatory network of **Figure 6D**. **D**, The expression level of *TaDOF17-B1*, *TaMYB30-A1*, *WFZP-D1*, *TaAP2-39-D1*, *TaBAK1-D1* and *TaPILS7-A1* across spike development.

**Supplemental Figure 8.**
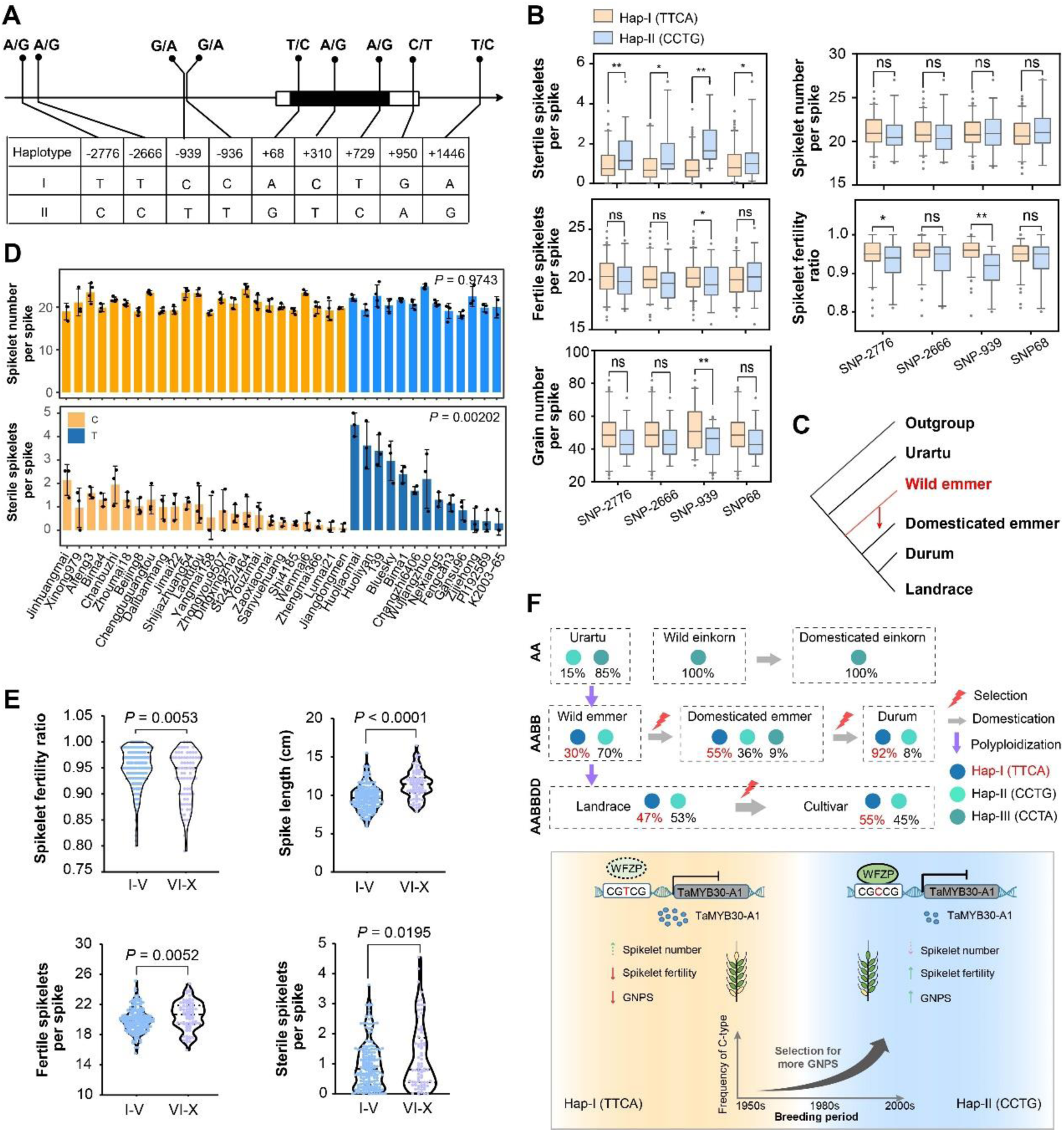
The expression regulation and phenotype analysis based on SNPs of *TaMYB30-A1.* **A,** Schematic diagram showing the polymorphism at each position for Hap-I and Hap-II in common wheat population. **B,** The comparison of SSPS and FSPS between haplotypes is defined by four significantly associated SNPs. Wilcoxon rank-sum test was used to determine the statistical significance; *, *P* < 0.05; **, *P* < 0.01. **C**, A phylogenetic topology showing the origin and transmission of haplotype Hap-TTCA in wheat and relatives. **D**, The SNS and SSPS of 37 cultivars with different haplotypes of *TaMYB30-A1.* Cultivars with C allele and T allele were marked in yellow and blue, respectively. The SNS and SSPS were shown in bar with data from each environment in a dot, and significant difference of cultivars between two haplotypes was determined by Student’s *t*-test. **E**, Zones I-V have higher spikelet fertility ratio but smaller SL, FSPS and SSPS than zones VI-X. Wilcoxon rank-sum test was used to determine the statistical significance between two groups. **F**, Schematic illustration of the selection of *TaMYB30-A1* excellent haplotype during domestication (upper panel) and breeding (bottom panel) process. Hap-I emerged in tetraploid and underwent human selection during the domestication of tetraploid and hexaploid wheat. The Hap-I with C-allele of *TaMYB30-A* promoter region within WFZP binding motif was selected during the breeding process in China over past 40 years. The haplotype (different colored circles) and their frequency (with Hap-I marked in red) are shown for each species. The purple vertical arrows indicate the phylogenetic relatedness between taxa, and gray horizontal arrows for domestication process. The natural/artificial selection of *TaMYB30-A1* excellent haplotype during domestication or improvement is indicated with a red lightning symbol. Blunt (┴) in the figure represents inhibition effects; The intensities of inhibition effects are indicated by the thickness of the arrows. The fertile and sterile spikelets are indicated in green and yellow, respectively.

**Supplemental Figure 9.**
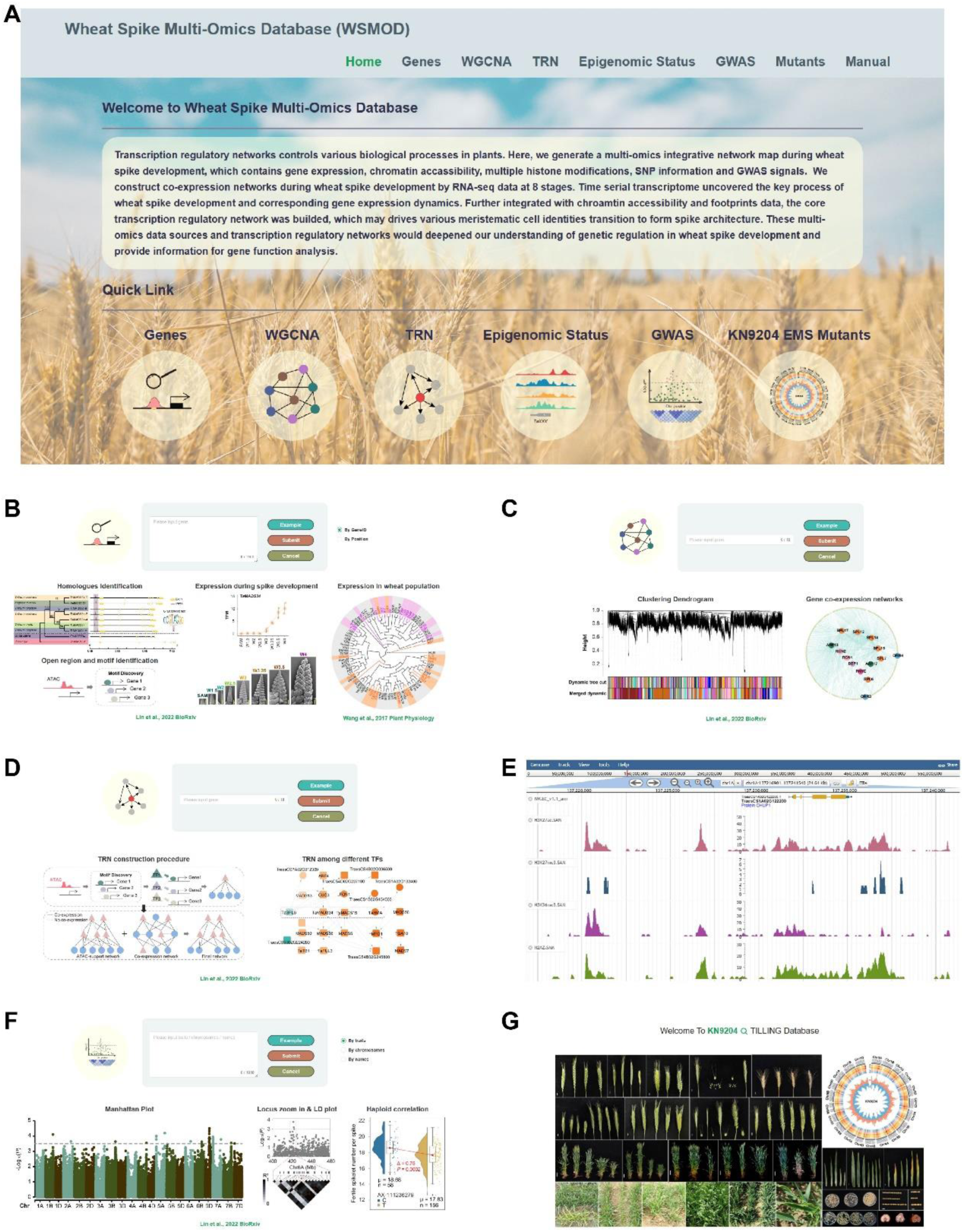
The overview of Wheat Spike Multi-Omics Database (WSMOD). **A,** The homepage of WSMOD. **B-G,** The six primary functional modules that comprise WSMOD. (**B**) Search by gene ID function provides gene expression patterns, WGCNA modules, orthologs in rice and *Arabidopsis thaliana*, open chromatin regions and footprints, QTL information. (**C**) Co-expression network function provides gene-gene co-expression relationships. (**D**) The wheat spike TRN function provides information of genes upstream TFs, TF binding sites and motifs, as well as downstream target genes. (**E**) Epigenomic status function offers visualization of transcription, chromatin accessibility, various Histone modifications, variations and QTL site. (**F**) GWAS function provides spike-related QTL regions and gene information in regions. (**G**) KN9204 EMS Mutants function provides a quick link to KN9204 TILLING Database in which users can conveniently query and order the EMS mutants of the interested gene.

## Notes

### Competing Interest Statement

The authors have declared no competing interest.

### Summary of Updates

Updated and supplemented some data and results

